# Determination of oligomeric organization of membrane proteins from native membranes at nanoscale-spatial and single-molecule resolution

**DOI:** 10.1101/2023.02.19.529138

**Authors:** Gerard Walker, Caroline Brown, Xiangyu Ge, Shailesh Kumar, Mandar D. Muzumdar, Kallol Gupta, Moitrayee Bhattacharyya

## Abstract

The oligomeric organization of membrane proteins in native cell membranes is a critical regulator of their function. High-resolution quantitative measurements of oligomeric assemblies and how they change under different conditions are indispensable to the understanding of membrane protein biology. We report a single-molecule imaging technique (Native-nanoBleach) to determine the oligomeric distribution of membrane proteins directly from native membranes at an effective spatial resolution of ∼10 nm. We achieved this by capturing target membrane proteins in “native nanodiscs” with their proximal native membrane environment using amphipathic copolymers. We established this method using structurally and functionally diverse membrane proteins with well-established stoichiometries. We then applied Native-nanoBleach to quantify the oligomerization status of a receptor tyrosine kinase (TrkA) and a small GTPase (KRas) under conditions of growth-factor binding or oncogenic mutations, respectively. Native-nanoBleach provides a sensitive, single-molecule platform to quantify membrane protein oligomeric distributions in native membranes at an unprecedented spatial resolution.

## Introduction

The cell membrane provides a dynamic platform that controls the organization, topology, and bioactivity of membrane proteins. The oligomeric organization of membrane proteins in their native environment plays a critical role in regulating their physiological function and serves to coordinate crosstalk between cells and their environment. A quantitative understanding of membrane protein oligomerization requires analytical tools to identify and monitor the distribution of membrane protein oligomeric assemblies directly from the native cell membrane environment at simultaneously high molecular and spatial resolution^1–3^. Advances in light microscopy have provided insights into the organization of macromolecular assemblies in cells and complex tissues^4–7^. Yet determining the precise oligomeric status for membrane proteins and structural details underlying these assemblies remains a challenging problem. Membrane protein oligomeric states are commonly characterized in cells or on membrane-mimetic platforms by diffusion and fluctuation measurements using fluorescence correlation spectroscopy (FCS)^8,9^, subunit counting using photobleaching step analysis^10,11^, single-molecule localization microscopy (SMLM)^4,5,12–14^, and single-particle tracking^15–17^. Cell-based readouts preserve the native membrane environment but often lack sufficient spatial and molecular resolution to distinguish genuine oligomeric assemblies from spatially-proximal colocalization^4,5,18,19^. Membrane-mimetic platforms, on the other hand, offer molecular resolution but fail to mimic the complexity of mammalian membrane environments, which may influence the oligomeric organization of embedded proteins^1,2,20–22^.

The main challenge lies in obtaining high lateral spatial resolution and quantitative molecular resolution simultaneously while preserving the surrounding native-membrane environment. The diffraction limit of light restricts the capacity to resolve particles that are closer than 200 nm in space using light microscopy^23^. This prevents unambiguous determination of stoichiometry due to the inability to distinguish protein subunits that form a *bona fide* oligomeric interface from those that are simply in close proximity on the membrane^23^. Another significant challenge with inferring oligomeric distributions using single-molecule imaging is obtaining single-molecule dilution of proteins embedded in or bound to membranes. For soluble proteins, this is easily done through dilution. However, the presence of the lipid bilayer precludes translation of such strategies to membrane proteins without disrupting their native environment through the use of membrane mimetics. Genetic and other approaches can lower protein concentrations on cell membranes for single-molecule detection^11,24,25^. But this could alter physiological oligomeric distributions if the protein concentration is reduced below the required threshold for assembly^26^. Under-labeling of membrane proteins with fluorescent moieties provides an alternate approach to achieving single-molecule density on native membranes, but ignores the participation of unlabeled subunits in observed oligomeric assemblies^25^. Therefore, new methods that afford single-molecule resolution of membrane proteins within their native membrane environment at endogenous, physiologically-relevant (high or low) expression levels are needed.

Addressing these collective challenges, we present a total internal reflection fluorescence (TIRF) microscopy-based single-molecule photobleaching step analysis technique that directly detects and quantifies the oligomeric distribution of membrane proteins in 8 - 12 nm circular patches of native membranes or “native nanodiscs”. Our approach isolates these native nanodiscs by detergent-free solubilization of cell membranes using amphipathic copolymers (such as styrene maleic acid (SMA) or styrene acrylic acid (AASTY), *Fig. 1a*)^27–30^. The solubilized GFP-tagged membrane proteins are surrounded by an annular ring of native lipids and neighboring proteins held together by these amphipathic copolymers. Native nanodiscs encapsulating target membrane proteins are soluble in aqueous buffer and can be further diluted before immobilizing on functionalized glass substrates to achieve single-nanodisc spot density (*Fig. 1d-e*). We then perform single-molecule photobleaching step analysis on each native nanodisc-membrane protein complex to identify their oligomeric states (*Fig. 1f*). We name this method Native-nanoBleach.

**Figure 1:**
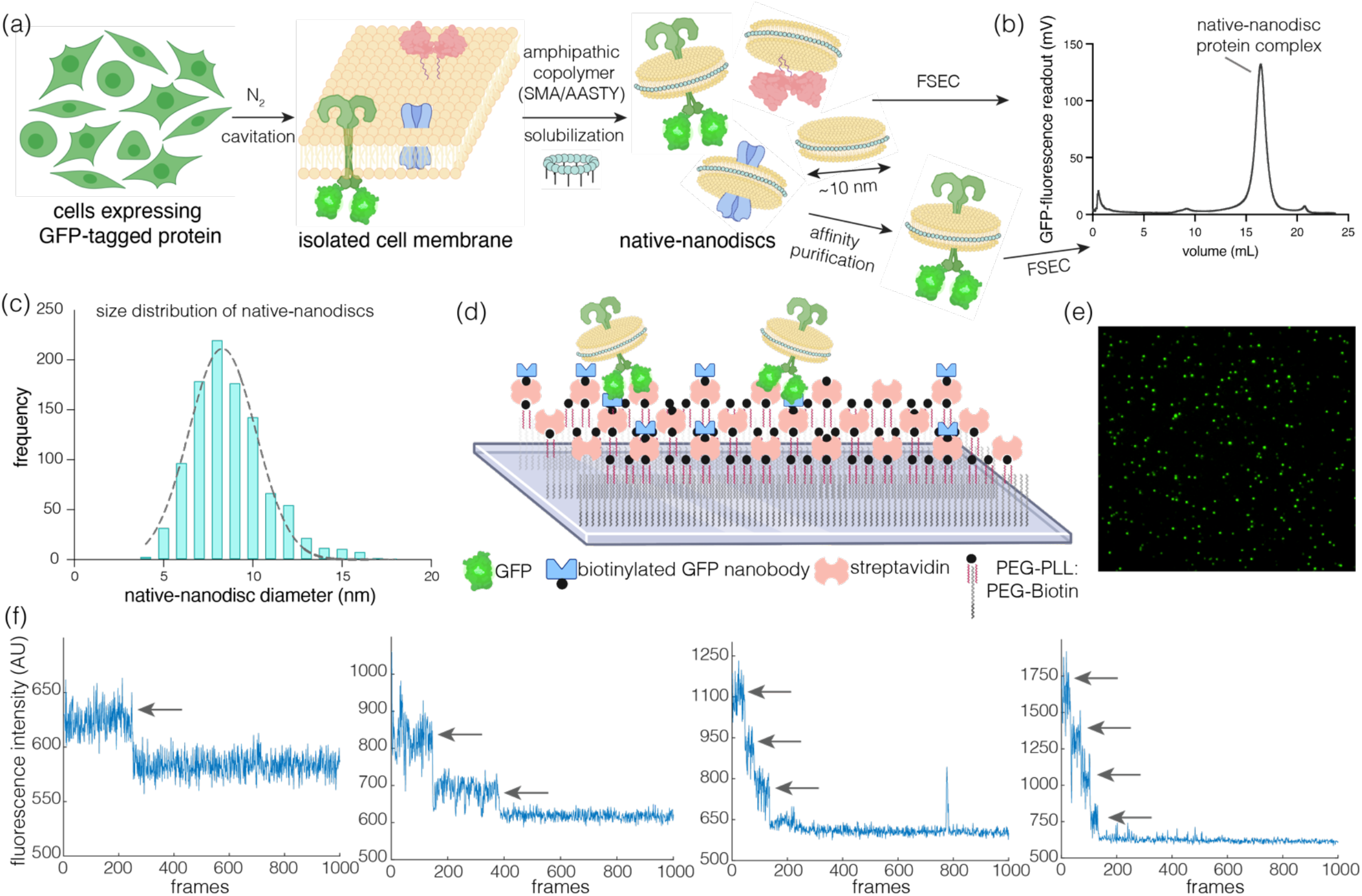
Native-nanoBleach approach to detect the oligomeric distribution of membrane proteins in native membrane environment. (a) Schematic diagram of the workflow for native-nanoBleach sample preparation. Membranes isolated from cells expressing GFP-tagged membrane proteins were solubilized with amphipathic copolymers to generate native nanodiscs. Native nanodiscs undergo either fluorescence detection size exclusion chromatography (FSEC), or affinity purification followed by FSEC. (b) Representative FSEC trace is shown for a native nanodisc encapsulated membrane protein (KRas WT) monitored for GFP fluorescence at 488 nm. (c) Size distribution of native nanodiscs by quantification of negative-stain EM images. Histogram shows the distribution of native nanodisc-KRas diameters (in nm) measured over ∼1000 particles. (d) Depiction of the setup for single-molecule TIRF-based photobleaching step analysis. The cartoon details the functionalization of glass slides (substrates) to display a GFP nanobody trap for capturing GFP-tagged membrane proteins within native nanodiscs. (e) Representative single-molecule TIRF image where each green spot represents a single native nanodisc encapsulated protein. (f) Representative photobleaching traces showing decrease in GFP intensity over time (9.3 frames/sec) for proteins bearing one, two, three, and four mature GFP subunits. The steps in each trace are highlighted using arrows.

Native-nanoBleach analysis has three key capabilities. First, the effective lateral spatial resolution is imposed by the diameter of native nanodiscs (8 - 12 nm), restricting imaging to within a region ∼25-fold smaller than the diffraction limit of light^5,23^. In our method, we count membrane protein subunits within each native nanodisc using photobleaching step analysis to determine oligomeric distributions. This analysis offers an unprecedented effective spatial (lateral) resolution of ∼10 nm with commercially available TIRF microscopes while maintaining the surrounding native environment around membrane proteins^5,23^. Second, each isolated native nanodisc-membrane protein complex represents a single particle in our photobleaching analysis. This eliminates the need for any prior adjustment to protein expression levels or labeling optimization on cell membranes to achieve single-molecule density as required for cell-based readouts. This allows interrogation of membrane protein oligomeric states at concentrations comparable to their endogenous expression levels. Finally, Native-nanoBleach does not require cell fixation as with many super-resolution imaging techniques for studying oligomerization^4,5,12^, thereby better preserving the native membrane environment.

Here, we have established the Native-nanoBleach approach by studying functionally and structurally diverse membrane proteins with well-established oligomeric states. We then applied our method to determine the oligomeric organization of selected members of the receptor tyrosine kinase (RTK)-Ras-MAPK signaling pathway, which is critical to cell growth, differentiation, and survival. We specifically studied the stoichiometry of the nerve growth factor (NGF) receptor TrkA^31,32^ and KRas4B (called KRas hereafter)^33,34^ under conditions of growth factor stimulation and oncogenic mutations, respectively. Using Native-nanoBleach, we analyzed the stoichiometry of TrkA before and after NGF binding to determine the growth-factor induced changes in TrkA oligomerization. We also demonstrate the presence of KRas dimers in native nanodiscs extracted from Expi293 cells overexpressing KRas as well as pancreatic ductal adenocarcinoma (PDAC) cells that express physiological levels of KRas^35^. We further show that dimerization is increased in oncogenic KRas variants. Since previous data on reconstituted supported bilayers showed that KRas has no intrinsic dimerization properties^8^, these results highlight the critical contribution of the native membrane environment in KRas dimerization.

With advances in CRISPR-Cas9 gene-editing technologies, it has become routine to attach fluorescent tags to endogenous proteins in animal models or patient-derived cells^36–38^. This portends broad applications of Native-nanoBleach analysis to determine the oligomeric organizations of endogenously-tagged membrane proteins in their native membrane environments, under various physiologically and clinically-relevant conditions, at ∼10 nm spatial and single-molecule resolution.

## Results

### A single-molecule assay (Native-nanoBleach) to determine membrane protein oligomeric organization in native membrane environment

Our approach uses commercially available amphipathic copolymers (SMA or AASTY) to isolate native nanodiscs (or circular patches) of unperturbed cell-membranes in a detergent-free extraction strategy (*Fig. 1a-b*)^27–30,39^. Such excision of isolated cell membranes to generate native nanodiscs enables isolation of membrane proteins in annular rings of neighboring native-lipids and proteins held together by the copolymers, preserving their corresponding proximal membrane environments. We purified native nanodiscs bearing fluorescently-tagged membrane proteins of interest by a combination of affinity and fluorescence-detection size exclusion chromatography (FSEC). We used the FSEC profiles to estimate the homogeneity and quality of the native nanodisc-target membrane protein complexes, monitoring the emission of GFP (*Fig. 1b*)^40^. The GFP-positive non-void volume fractions corresponding to native nanodisc encapsulated target membrane proteins were collected and used for downstream analysis. We used negative-stain electron microscopy (negative stain EM) to measure the size distribution of the native nanodiscs obtained using SMA or AASTY, which displayed a narrow distribution with a mean of ∼8.5 nm (SD 2 nm) and ∼12 nm (SD 3 nm), respectively, across diverse samples and over thousands of particles measured (*Fig. 1c, Supplementary Fig. 1*).

We designed a TIRF-based single-molecule photobleaching step analysis (Native-nanoBleach) to determine the oligomeric distribution of membrane proteins in individual native nanodiscs at a lateral spatial resolution of ∼10 nm. Our method relies on capturing well-separated single particles of native nanodisc GFP-tagged membrane protein complexes on glass substrates functionalized with biotinylated GFP-nanobody (*Fig. 1d, Supplementary Fig. 2*). Each GFP-fluorescent spot in our TIRF images represents a single native nanodisc membrane protein complex (*Fig. 1e*). Substrate immobilization can be customized to support other affinity or antibody traps based on the specific protein tags used. The single-molecule detection ensures high sensitivity, allowing application of this experimental platform to study low abundance proteins at endogenous expression levels. The photobleaching data was acquired by continuously exposing an area on the glass substrate to laser in TIRF mode, and movies of 1500-2000 frames were collected at ∼9.3 frames/second. These data were analyzed by plotting the time-dependent decrease in GFP fluorescence intensity for each spot (*Fig. 1f*). For each sample, discrete photobleaching steps were counted for a total of 3000-4000 particles from at least three biological replicates. These data are represented by a distribution ranging from 1 to 4 steps.

We obtained the oligomeric landscape of membrane proteins from this experimentally obtained step-distribution. The number of photobleaching steps underrepresents the number of subunits within an oligomer due to ∼70% maturation efficiency of GFP^11,41^. For example, 1-step photobleaching may arise from monomers but also from some dimers with only one subunit carrying a mature, fluorescent GFP. We accounted for the dark GFP state as follows. We first obtained the theoretical probability distribution of how many subunits are GFP fluorescent within a given oligomeric state (from monomer-tetramer), assuming GFP maturation as 70%. For proteins exhibiting a single stoichiometry (either monomer, dimer, trimer, or tetramer), we can directly match the experimental step distribution with one of these theoretical step distributions to identify the specific oligomeric state (*Fig. 2a*). For proteins with oligomeric distributions spanning across monomer to tetramer, we further calculated the relative proportions of monomers, dimers, trimers, and tetramers whose combined theoretical distribution best fit the experimentally-obtained step distribution (*Fig. 2b-c*). A detailed description of these calculations is provided in the Online Methods section.

**Figure 2:**
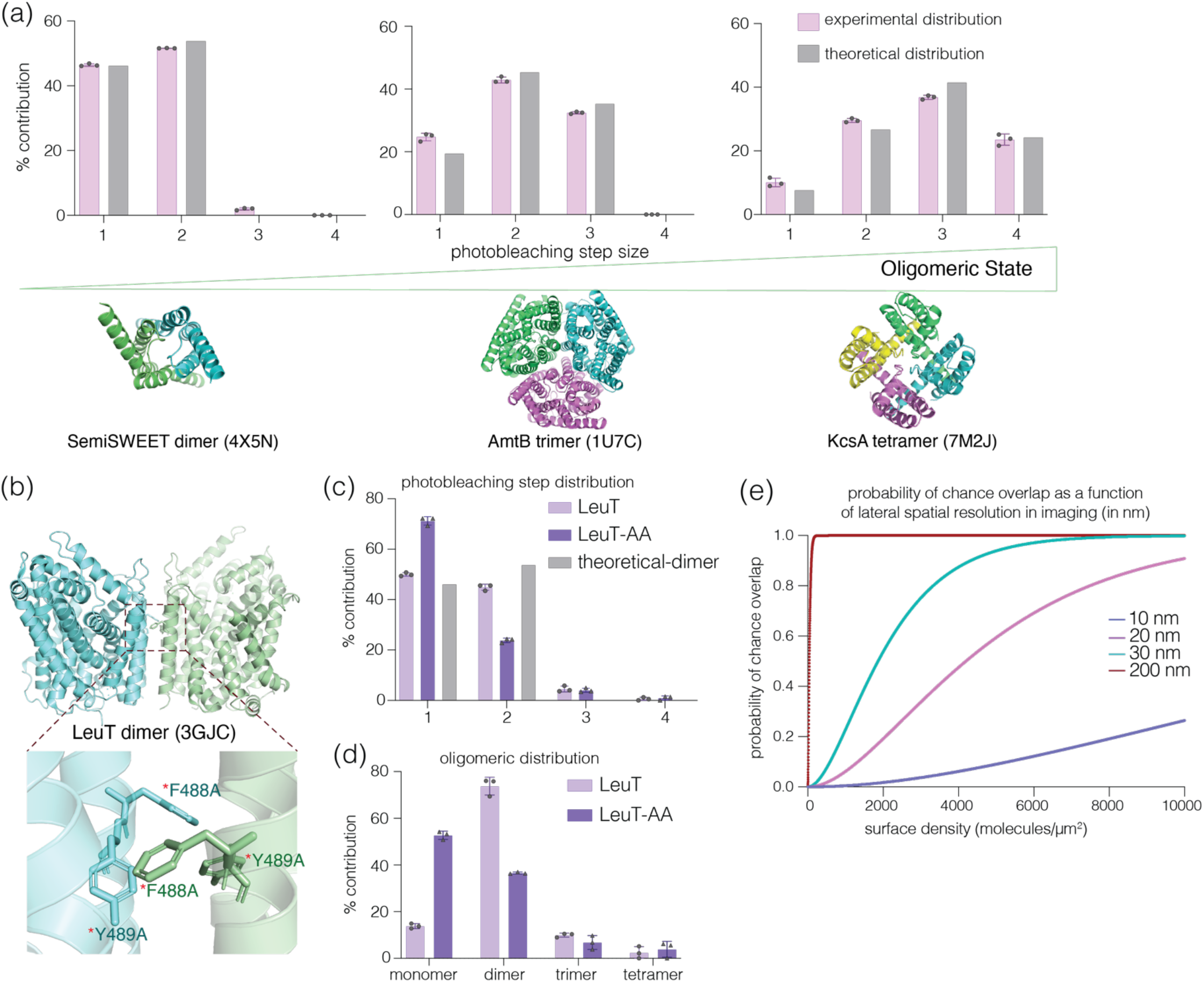
Establishing the native-nanoBleach analysis using membrane proteins with well-established oligomeric states. (a) GFP-photobleaching step distribution of SemiSWEET (dimer), AmtB (trimer), KcsA (tetramer). Each panel compares the experimentally-observed distribution of 1-4 steps with theoretical step-distribution corresponding to dimer, trimer and tetramer, respectively, assuming 70% maturation efficiency of GFP. (b) Cartoon depiction of the dimeric structure of LeuT (PDB_ID: 3GJC). The zoomed panel highlights the mutated residues in the LeuT-AA mutant, F488 and Y489, which stabilize the dimeric interface through pi-stacking interactions. (c) Comparison of experimentally observed step-distribution of LeuT and LeuT-AA with the theoretical distribution corresponding to a dimer. (d) Oligomeric distribution of LeuT and the dimer interface mutant (AA) that matches the step distribution in (c), as calculated considering GFP maturation to be 70%. All photobleaching step-analysis shown in (a, c-d) is from a total of about 3000-4000 spots over at least three biological replicates. Data shown as mean ± sd. (e) Plot showing the theoretical probability of two protein subunits randomly colocalizing in a 10 nm X 10 nm area for a range of membrane protein surface densities (or membrane protein expression levels) in molecules/µm^2^.

### Establishing the Native-nanoBleach approach using a diverse set of membrane proteins with distinct oligomeric states

To establish the broad applicability of our platform, we chose a diverse set of membrane proteins, expressed in *E. coli* as well as various mammalian cells, with stoichiometries ranging from monomer to tetramer, containing 1-12 transmembrane helices per subunit, and representing diverse biological functions (*Table S1*). Among these, we used four integral membrane proteins with well-defined stoichiometries to demonstrate the capabilities of Native-nanoBleach and validate the method – the sugar transporter SemiSWEET (dimer)^42^, the ammonium transporter AmtB (trimer)^43,44^, the potassium ion channel KcsA (tetramer)^45,46^, and the amino-acid transporter LeuT (monomer-dimer mixed populations)^47^. Each protein was tagged with a monomeric variant of enhanced GFP^48,49^ (referred to as GFP hereafter) at the C-terminus^48,49^. We solubilized these proteins using amphipathic copolymers and performed FSEC to remove any aggregates (*Supplementary Fig. 3*). We captured the GFP-positive non-void fractions for each sample on functionalized substrates at single-molecule density and analyzed stoichiometries using native-nanoBleach.

The experimentally obtained photobleaching step distributions for SemiSWEET, AmtB, and KcsA closely matched the theoretical distribution for a dimeric, trimeric, and tetrameric protein, respectively (*Fig. 2a*). We further demonstrated, using LeuT as an example, the ability of our platform to identify perturbation-induced redistribution of oligomeric populations for membrane proteins that exist in an equilibrium of different oligomeric states. We showed that LeuT is predominantly dimeric (75%) with a small fraction of monomers (14%) (*Fig. 2b-d*). We next introduced mutations at the LeuT dimer interface (F488A and Y489A) based on its reported crystal structure (PDB_id: 2A65)^50^. The alteration of these residues ablates critical pi-stacking interactions and is predicted to weaken the LeuT dimeric interface (*Fig. 2b*). This mutant (LeuT-AA) exhibited an increased monomeric population (53%) and decreased dimers (36%), as expected. These results are in good agreement with previous native mass spectrometry analysis of LeuT oligomerization^47^.

Together, our data establish the Native-nanoBleach platform for deciphering the oligomeric organization of diverse classes of membrane proteins in their native environment at ∼10 nm lateral spatial resolution. Further, as shown with LeuT, the platform enables us to quantitatively illustrate how mutations dynamically regulate such assemblies in native membranes, which can be extended to study effects of ligand or drug binding and environmental perturbations on oligomeric distributions. Based on the close agreement between the oligomeric status determined from Native-nanoBleach and those reported in the literature for these membrane proteins, we established that the amphipathic copolymers in native nanodiscs do not interfere with the determination of oligomeric distribution (*Fig. 2a-d*). We also established that the fraction of multi-step photobleaching events occurring due to coincidental colocalization of two or more protein subunits within a nanodisc or two or more nanodiscs in the same spot on functionalized substrates is insignificant.

To understand the correlation between expression levels of membrane proteins and chance colocalization of individual subunits within native nanodiscs, we calculated the theoretical probability of two or more protein subunits to randomly colocalize within a 10 × 10 nm^2^ area as a function of a wide range of membrane protein surface densities (in molecules/*μ*m^2^) (*Fig. 2e*, see Online Methods for details). We estimated that, under conditions where the membrane protein surface density is less than 1000 molecules/*μ*m^2^, there is less than 0.5% probability of coincidental colocalization of protein subunits within 10 × 10 nm^2^ on membranes. At a spatial resolution of 20, 30, or 200 nm, the probability for chance overlap increases to 6%, 22%, and 100%, respectively, for a surface density of 1000 molecules/*μ*m^2^ (*Fig. 2e*). These data confirm the importance of lateral spatial resolution in determining oligomeric status of membrane proteins at varying levels of expression. The estimated surface density of various membrane proteins, as described in the literature, rarely exceeds 1000 molecules/*μ*m^2^ (for example, even EGFR overexpressing A431 cancer cells show surface density of 636 molecules/*μ*m^2^)^51,52^. Therefore, the Native-nanoBleach approach, with its ∼10 nm effective lateral spatial resolution, is capable of discriminating chance overlap from *bona fide* protein oligomerization over a wide range of surface expression levels.

### Application of Native-nanoBleach to determine the stoichiometry of the nerve growth factor receptor TrkA on native membranes

We next applied Native-nanoBleach to study ligand-induced changes in the oligomeric distribution of TrkA, an RTK that plays crucial roles in neuronal development and maintenance in response to growth factors (specifically neurotrophins such as NGF)^31^. The canonical mechanism of RTK activation involves the transition from unliganded monomers to growth factor-induced dimers or higher-order oligomers^32^. This promotes intracellular kinase activation and subsequent autophosphorylation that regulates various downstream signaling pathways. We analyzed the TrkA oligomeric assemblies and their relative populations in the presence or absence of NGF in native nanodiscs isolated from SH-SY5Y neuroblastoma cells (*Fig. 3a*). We selected the SH-SY5Y expression system as these neuroblastoma cells display neurite outgrowth in response to NGF, providing a physiologically relevant cellular backdrop for our analysis^53,54^. In addition, expression of TrkA in SH-SY5Y cells led to relatively lower basal levels of autophosphorylation (pY490) as compared to overexpression in Expi293 cells and shows increase in pY490 levels upon NGF treatment (*Supplementary Fig. 4*).

**Figure 3:**
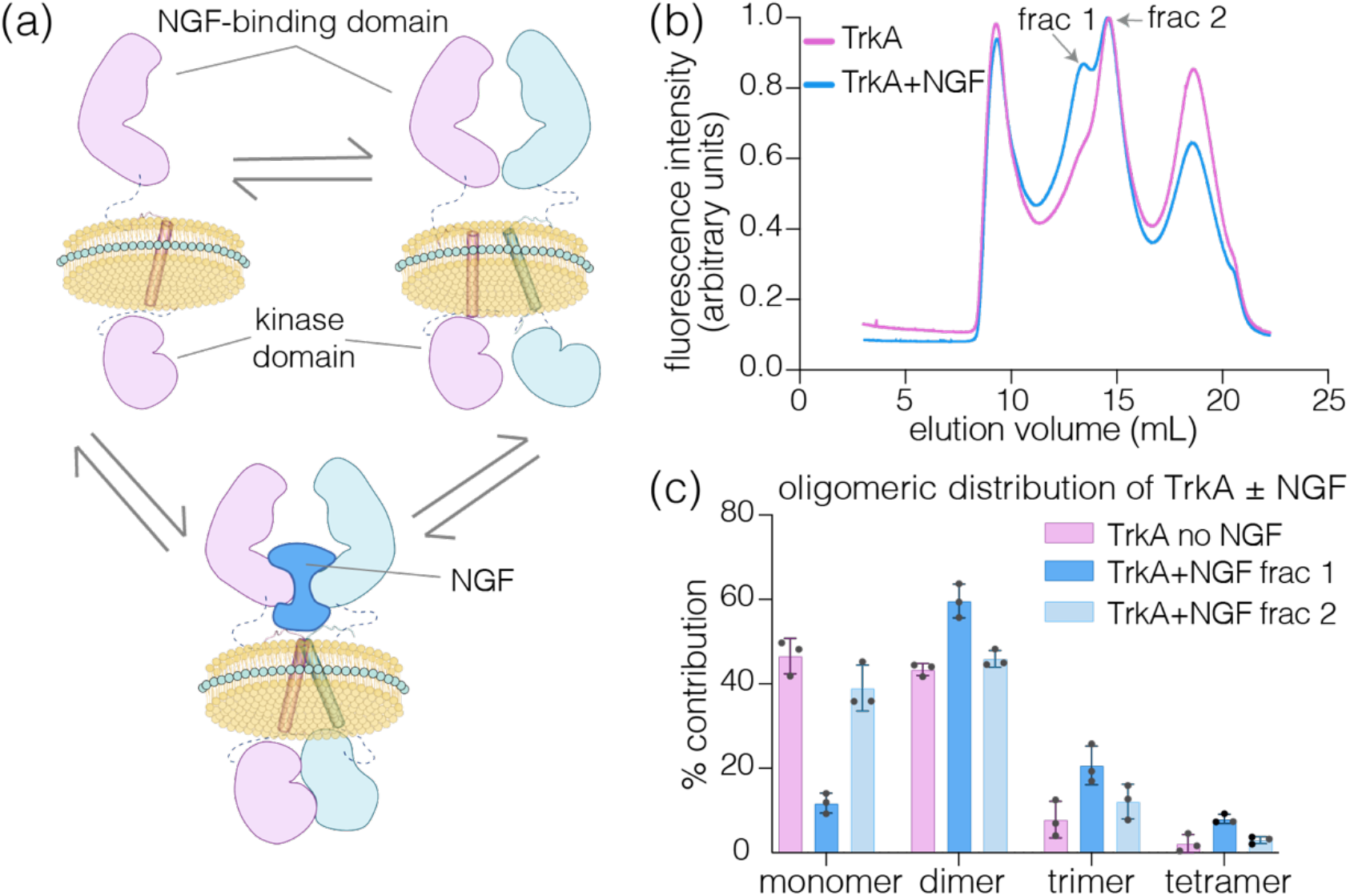
Oligomeric distribution of TrkA in the presence and absence of NGF within native nanodiscs isolated from SH-SY5Y cells. (a) Schematic showing various oligomeric states of TrkA, including monomer, ligand-independent dimers, and NGF-mediated dimers (higher-order oligomers are not shown for clarity). (b) FSEC profile of native nanodisc encapsulated TrkA isolated from cells treated with NGF or untreated cells monitoring C-terminal GFP. Treatment with NGF leads to the formation of a left peak (fraction 1) in addition to a peak corresponding to that seen in the untreated sample (fraction 2), indicating the emergence of higher order oligomers. The FSEC traces are normalized to have the same area under the curve. (c) Native-nanobleach analysis of frac 1 and 2 from untreated and NGF-stimulated samples shows TrkA oligomeric distribution under these conditions. Data in (c) are from a total of about 3000-4000 spots over at least three biological replicates represented as mean ± sd.

Our Native-nanoBleach analysis revealed TrkA to exist as a mix of 46% monomer, 44% dimer, and 10% higher-order oligomers in the absence of NGF (*Fig. 3b-c, Supplementary Fig. 5*). After treatment of TrkA expressing SH-SY5Y cells with 100 ng/ml of NGF for 15 min, we measured growth-factor induced redistribution of TrkA oligomers by FSEC and Native-nanoBleach. The FSEC profile shows a new peak at a lower elution volume, indicating an increased proportion of higher order oligomers in addition to a peak corresponding to untreated TrkA (fractions 1 and 2, respectively; *Fig. 3b*). Fraction 1 predominantly exhibits TrkA dimers (∼60%) and higher-order oligomers (∼30%), whereas fraction 2 shows similar distribution to untreated TrkA (*Fig. 3c*). Our results are in agreement with previous studies on RTKs that show a departure from their canonical mechanism of activation, including ligand-independent dimerization of TrkA^55–60^.

Alterations in the oligomeric status for RTKs have been associated with pathological mutations, truncations, and oncogenic fusions, which in turn regulate the activation of these transmembrane kinases on the plasma membrane^32,61^. Using TrkA as an example, we have established Native-nanoBleach as an experimental platform that can be used to quantify the ligand-independent and ligand-induced dimerization and higher-order oligomerization of RTKs (58 members categorized into 20 subfamilies) on native membranes under various physiological and pathological conditions.

### Application of Native-nanoBleach to determine the oligomeric status of KRas and its oncogenic mutants in their native membrane environment

Downstream of RTKs, the small oncogenic GTPase Ras forms nanoclusters on the plasma membrane, acting as a regulatory hub for membrane-localized signaling pathways^62,63^. Whether the building blocks of these nanoclusters are formed by Ras dimers/oligomers or simply spatially-concentrated individual Ras subunits has remained the subject of intense investigation^8,14,62,64^. To answer this question, we have to first address whether KRas forms dimers/oligomers when bound to native membranes. Native-nanoBleach analysis addresses this question by interrogating the native nanodisc-KRas complex at ∼10 nm spatial and single-molecule resolution in the context of the native membrane environment. As discussed in *Fig. 2e*, KRas subunits within 10 nm of each other are likely due to specific interactions (either direct KRas interactions or indirect interactions mediated by native lipids or neighboring proteins) for a wide range of KRas concentrations.

We isolated and purified native nanodiscs containing KRas from Expi293 cell membranes expressing 6X-His-GFP-tagged KRas and its oncogenic mutants, G12D and Q61H^34^ (*Supplementary Fig. 6a*). Native nanodisc-KRas showed 54% monomer, 40% dimer, and ∼5% higher-order oligomers as calculated from the experimental photobleaching step distributions (*Fig. 4a, Supplementary Fig. 7a*). Introduction of known oncogenic mutations in KRas led to a significant increase in dimers (53% and 48%) and higher-order oligomers (17% and 14%) for G12D and Q61H, respectively, with a concomitant decrease in monomers (*Fig. 4a*). We also demonstrated that the oligomeric organization remained identical between native nanodisc KRas-G12D purified using SMA or AASTY despite the slightly larger size of the AASTY-nanodiscs (*Supplementary Fig. 8*). This result provides additional evidence against possible coincidental localization of KRas subunits on the membrane, as that would lead to an increase in dimers/higher-order oligomers with increasing nanodisc size.

**Figure 4:**
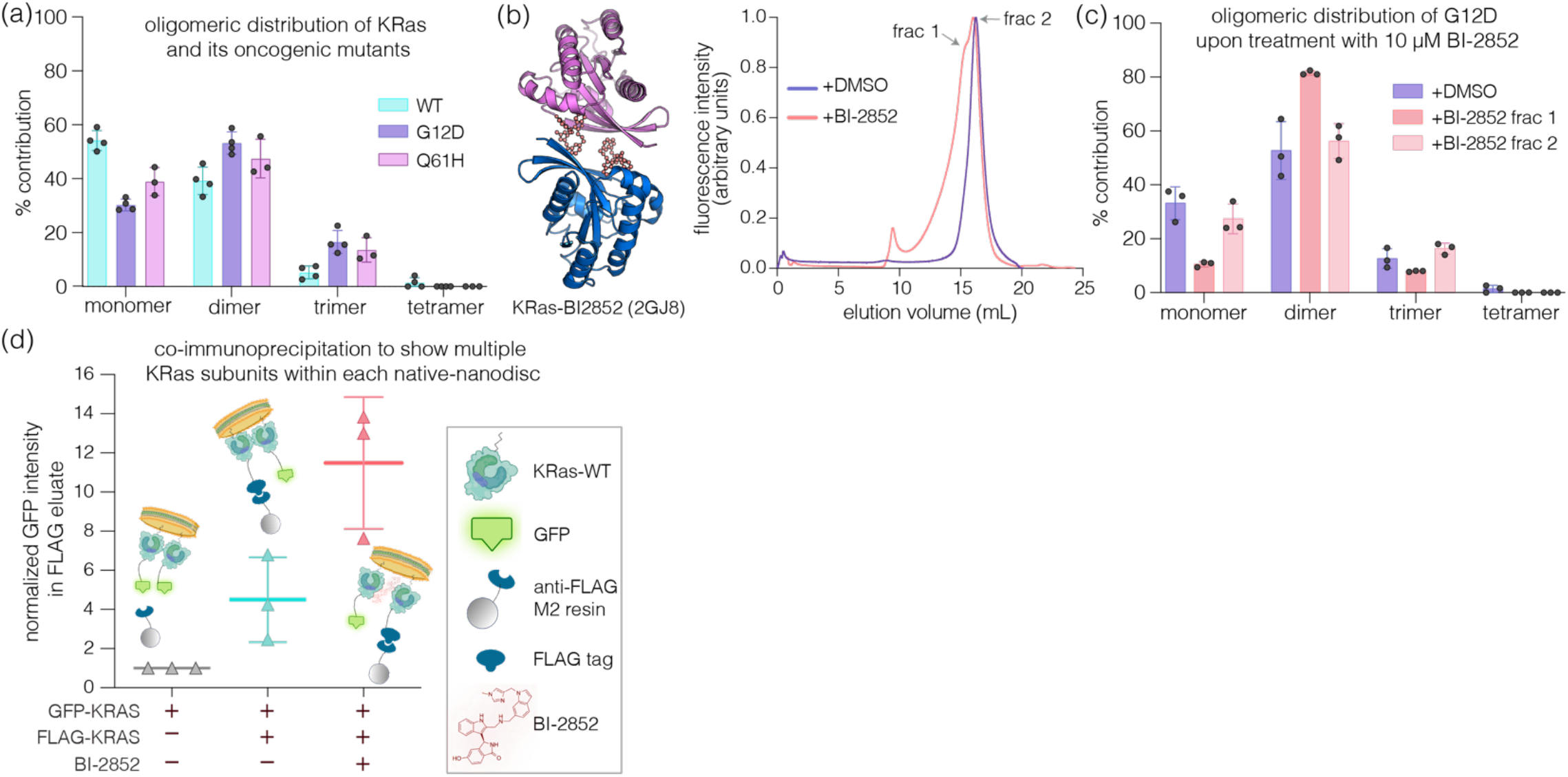
Oligomeric distribution of KRas and its oncogenic mutants within native nanodiscs isolated from Expi293 cells. (a) Native-nanoBleach analysis of KRas WT, G12D, and Q61H encapsulated in native nanodiscs. (b) FSEC traces for native nanodiscs encapsulating KRas-G12D isolated from cells treated with BI-2852 (in DMSO) or DMSO alone. BI-2852 treatment induces appearance of a left-shifted peak on the FSEC trace (fraction 1) in addition to a peak a peak corresponding to that seen in only DMSO-treated cells (fraction 2). The FSEC traces are normalized to have the same area under the curve. (c) Native-nanobleach analyses of fractions 1 and 2 reveal BI-2852-induced enhancement in KRas dimerization. The left-shifted frac 1 exhibits high proportion of dimers (>80%), acting as a positive control for our native-nanoBleach analysis, while frac 2 shows an almost identical distribution to that observed in untreated KRas-G12D. Data shown in (a, c) are from a total of about 3000-4000 spots over at least three biological replicates represented as mean ± sd. (d) Quantification of western blots from co-immunoprecipitation studies to confirm presence of multiple KRas subunits within each native-nanodisc. Expi293 cells transiently transfected with combinations of GFP-KRas and FLAG-KRas (+/-BI-2852) were purified using anti-FLAG M2 resin. Band densitometry (using Fiji) was used to quantify eluted GFP-KRas, normalized with respect to the total protein loaded onto anti-FLAG resin (estimated by Na/K-ATPase levels). The value obtained for the GFP-KRas only sample (negative control) is set to 1.0, accordingly adjusting the values for the other samples for ease of comparison.

To further confirm that these photobleaching alterations were indeed due to KRas dimerization, we repeated our analyses using BI-2852^65^, a drug proposed to induce non-functional KRas dimerization in structural studies^66^ (*Fig. 4b*). We treated Expi293 cells expressing KRas-G12D with 10 *μ*M of BI-2852 for 120 min and purified native nanodisc-KRas from these cell membranes. The FSEC profile for this sample shows an additional left-shoulder peak absent in solvent (DMSO)-treated samples (*Fig. 4b*). We captured the individual FSEC fractions on substrates and employed Native-nanoBleach. The shoulder peak corresponded to ∼80% KRas dimers, whereas the main peak showed the same oligomeric distribution as DMSO-treated samples (*Fig. 4c, Supplementary Fig. 7b*). These data also demonstrate the sensitivity of our method to identify changes in oligomeric distributions of membrane proteins upon small molecule binding, which can be extended to study interactions with antibodies/nanobodies/monobodies or effectors/substrates.

We further corroborated the presence of KRas dimers using an orthologous co-immunopurification approach in solution. (*Fig. 4d, Supplementary Fig. 9*). We transfected Expi293 cells with a 1:1 mixture of FLAG-tagged and GFP-tagged KRas followed by purification of native nanodiscs using anti-FLAG resin. We detected significant GFP-KRas in the fractions eluted from the FLAG-resin (∼5-fold higher than samples from cells expressing GFP-KRas alone which show negligible GFP signal in the eluate). Additionally, we treated these cells with BI-2852, which dramatically increased the GFP signal (∼10-fold) in the eluate and acts as a positive control (*Fig. 4d, Supplementary Fig. 9*). This observed colocalization of GFP-KRas within native nanodiscs encapsulating FLAG-KRas establishes the presence of multiple KRas subunits within each nanodisc.

Together, these data document the first detection of dimers and higher-order oligomers in purified samples of membrane bound KRas at ∼10 nm spatial resolution in the context of their proximal native lipidome and proteome. The requirement of the native membrane environment for detection of KRas dimers points to the importance of endogenous lipids and neighboring protein partners in aiding this oligomerization. Our data also show enhanced dimerization of KRas oncoproteins, implicating the importance of targeting the complex KRas dimer interface^67^ with small molecules and antibodies/monobodies/nanobodies that can be screened for using Native-nanoBleach.

### KRas dimerization is observed in PDAC cells stably expressing physiological levels of KRas and its oncogenic mutants

We next applied Native-nanoBleach to determine the oligomeric states of KRas variants at endogenous levels of expression in a clinically relevant cellular context. KRas mutations drive >90% of pancreatic cancers^33,68^, making PDAC an ideal model for studying KRas oligomeric organization in native membranes. We previously used CRISPR/Cas9 to knock out endogenous KRas from 8988T PDAC cells (NP10/PDAC^KRasless^ cells)^69^. We lentivirally reintroduced either GFP-KRas WT or its oncogenic variant G12V to these cells (*Fig. 5a*). These cells were sorted by flow cytometry to isolate and generate stable cell lines expressing KRas variants (called PDAC^GFP-KRas^ hereafter) at levels comparable or lower than endogenous KRas in a panel of PDAC cells (*Figs. 5a-b, Supplementary Fig. 10a*). Importantly, reconstitution of KRas restores morphology, proliferative kinetics, and anchorage-independent growth comparable to that of parental cells^69^.

**Figure 5:**
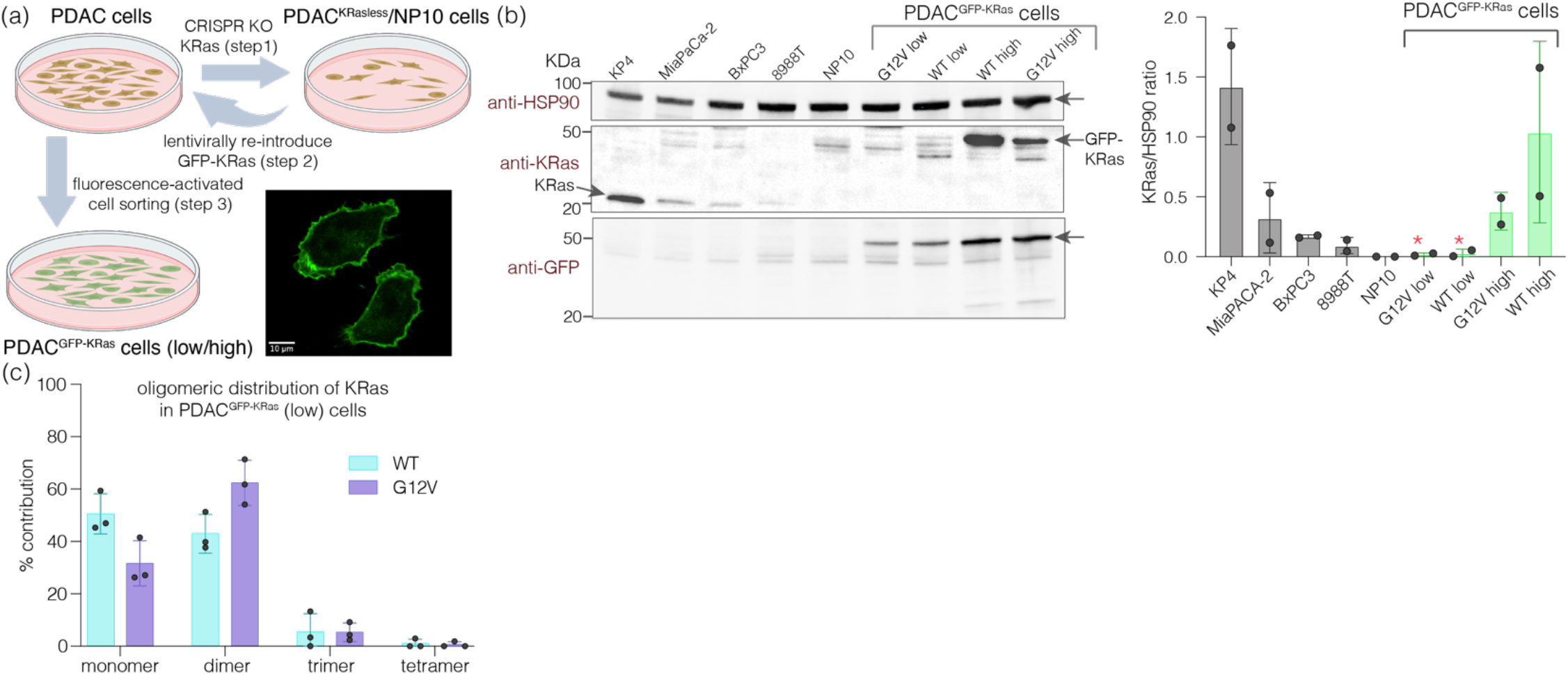
Oligomeric distribution of KRas and its oncogenic mutants within native nanodiscs isolated from PDAC cells. (a) Schematic depicting the generation of PDAC^GFP-KRas^ cells lacking endogenous KRas expression (NP10/PDAC^KRasless^ cells). (b) Western blot analysis comparing levels of KRas in stable PDAC^GFP-KRas^ cells expressing high or low levels of GFP-tagged KRas variants (as sorted by flow cytometry) with endogenous KRas levels in a panel of patient-derived PDAC cell lines (left). KRas expression levels in each cell line were quantified using band densitometry and normalized with respect to the amounts of HSP90 loading control using Fiji (right). The * indicates the cells that were selected for Native-nanoBleach analysis. (c) Native-nanoBleach analysis of KRas WT and G12V in native nanodiscs isolated from PDAC^GFP-KRas^ (low) cells. Data shown in (c) are from a total of about 3000-4000 spots over at least three biological replicates represented as mean ± sd.

We studied the oligomeric organization of KRas and its oncogenic mutant (G12V) in native nanodiscs purified from the PDAC^GFP-KRas^ (low) cells with comparable expression to parental 8988T cells (*Fig. 5a-b, Supplementary Fig. 6b*). These cells provide disease-relevant native membrane environment and a KRas-less background to express the GFP-tagged KRas variants, preventing undercounting of oligomeric states due to endogenous dark KRas subunits. WT KRas showed ∼50% monomer and 43% dimer (*Fig. 5c, Supplementary Fig. 7c*). KRas-G12V exhibited an increased proportion of dimers (62%) with a concomitant decrease in monomers (30%). Despite the 5-fold lower expression levels of KRas variants in PDAC^GFP-KRas^ (low) as compared to the expression in Expi293 cells (*Supplementary Fig. 10b-c*), the oligomeric distributions of KRas (WT and oncogenic mutants) in native nanodiscs isolated from these two cell lines were almost identical (*Figs. 4a, 5c)*.

A recent study has estimated that there are about 75,000 KRas4B molecules/cell in MiaPaca-2 cells^52^. Based on ∼10-fold lower expression of GFP-KRas in PDAC^GFP-KRas^ (low) cells as compared to MiaPaca-2 (*Fig. 5b*), we estimate there to be about 7500 KRas4B/cell in these stable lines. Assuming an approximate cellular diameter of 20 *μ*m and even with 100% localization of KRas to plasma membrane, the surface density of KRas in PDAC^GFP-KRas^ (low) cells is estimated to be about 5 molecules/*μ*m^2^. At this density, the probability of two KRas molecules coincidentally localizing within ∼10 nm of each other is essentially zero (see *Fig. 2e*).

Taken together, our data clearly show that membrane-bound KRas forms dimers at a wide range of protein concentrations when its native membrane environment is preserved and that oncogenic mutations enhance dimer formation. The precise molecular mechanisms by which native lipids and neighboring proteins in native nanodiscs regulate KRas dimerization warrant further investigation and will be addressed in future studies.

## Discussion

We have established the Native-nanoBleach method as a general experimental platform to analyze the oligomeric organization of diverse classes of membrane proteins (monotopic, bitopic, polytopic) in their native membrane environment (*Table S1*). Our method has an effective lateral spatial resolution of ∼10 nm that is imposed by the average diameter of native nanodiscs, which allows us to distinguish spatially-proximal protein subunits from those complexed by specific biophysical interactions. The theoretical probability of two protein subunits randomly colocalizing over a range of membrane protein surface densities clearly demonstrates the challenge in determining the oligomeric organization of membrane proteins on cell membranes using diffraction-limited light microscopy (*Fig. 2e*)^4,23^. Our calculations show that for a protein surface density of 50 molecules/*μ*m^2^, the probability of chance overlap at the diffraction limited spatial resolution of 200 nm (i.e., two subunits within 200 nm or less of each other) is ∼60% in contrast to less than 0.1% when measured at 10-30 nm spatial resolution. With the ∼10 nm effective spatial resolution of Native-nanoBleach, we can study up to a membrane protein surface density of 1000 molecules/*μ*m^2^ with less than 0.5% chance of coincidental protein overlap within the native nanodisc. This provides a large dynamic range of membrane protein concentrations while determining their oligomeric distributions on native membranes.

The preservation of the native membrane locale around target proteins directly accounts for the role of native lipids or proximal protein neighbors in supporting macromolecular assemblies on membranes, which is not possible using *in vitro* reconstituted systems. Additionally, the isolation of samples within native nanodiscs followed by single-molecule analysis allows highly sensitive detection even at low endogenous levels of membrane proteins, as illustrated by our study of KRas in PDAC cells. Native-nanoBleach can efficiently analyze membrane protein stoichiometry from cell membranes isolated from one 35 - 100 cm tissue culture plate, depending on expression level. This low sample requirement makes it possible to perform this analysis on cells and tissues that are difficult to obtain or culture under various stimulation and perturbation conditions.

We established Native-nanoBleach using membrane proteins with well-established stoichiometries, ranging from monomer to tetramer. Using LeuT, we demonstrated that our analysis is sensitive to perturbation (interfacial mutation) induced redistribution of oligomeric equilibrium. Having established the assay, we next applied Native-nanoBleach to understand ligand, small-molecule, or mutation-induced changes in the RTK-Ras-MAPK signaling axis, specifically for the NGF receptor tyrosine kinase, TrkA, and KRas. Our data indicate that native-nanoBleach can be applied widely to examine the impact of disease-relevant mutations, truncations, ligand/small molecule binding, and changes in membrane lipid composition on the oligomeric distribution of diverse membrane proteins. Native-nanoBleach can also be used to decipher the molecular mechanisms of membrane protein oligomerization by observing alterations in stoichiometry upon systematic mutational analysis.

This general technology can be advanced to further broaden the scope of its applications. Using in-house synthesized amphipathic co-polymers capable of systematically generating native nanodiscs of increasing diameter (6-30 nm) with Native-nanoBleach, we can obtain insights into macromolecular assemblies over increasing sized circular patches of the native cell membrane. Using SMA and AASTY, we have already achieved average disc sizes of ∼8.5 nm and ∼12 nm, respectively. Our results so far have shown no differences between using SMA or AASTY copolymers in Native-nanoBleach (*Supplementary Fig. 8*). In future studies, we will extend the repertoire of copolymers for optimal solubilization and stability of diverse membrane proteins.

Although here we have mainly focused on homodimerization, Native-nanoBleach can be extended to unravel the heterooligomeric organization of proteins on native membranes by judiciously labeling interacting partners. Combining this approach with microfluidics devices, we can further reduce the number of cells required for the determination of oligomeric status. In summary, this report showcases a single-molecule experimental platform to identify the oligomeric organization of membrane proteins in their native environment at an unprecedented lateral spatial resolution while highlighting the future developments that can build upon this method.

## Methods

### Preparation of plasmids

#### Bacterial expression constructs

SemiSWEET (*Vibrio sp*., Uniprot ID: F9RBV9), AmtB (*Escherichia coli*, Uniprot ID: P69681), LeuT (*Aquifex aeolicus*, Uniprot ID: O67854), and LeuT variants were cloned into the pET16b backbone with a C-terminal GFP tag (the pET16b-LeuT vector was a gift from Dr. Eric Gouaux, Oregon Health & Science University). All GFP coding sequences used in this study are that of monomeric enhanced GFP^48,49^, and referred to as GFP hereafter. KcsA (*Streptomyces lividans*, Uniprot ID: P0A334) was cloned into the pQE-60 expression vector with a C-terminal 3C-protease site followed by a GFP tag (pQE-60-KcsA was a gift from Dr. Crina Nimigean, Weill Cornell Medicine).

#### Mammalian expression constructs

Human KRas (Uniprot ID: P01116) and its variants were cloned into the pEG-BacMam expression vector with GFP and 8X-His tags at the N-terminus (Addgene, see *Table S2*). For FLAG-tagged KRAS expression constructs, a 3X-FLAG tag was cloned in place of the 8X-His-tag in the above plasmid. Human TrkA (Uniprot ID: P04629) was cloned into the pEG-BacMam expression vector with the GFP and 8X-His tags at the C-terminus (Addgene: *Table S2*). All constructs with large domain insertions and deletions were made using standard protocols for Gibson assembly (New England Biolabs, MA). All point mutants used were generated using the standard Quikchange protocols (Agilent Technologies, Santa Clara, CA). Expected construct sequences were confirmed using Sanger sequencing (Genewiz, Azenta).

#### Cloning of KRAS CRISPR sgRNA and GFP-KRAS lentiviral constructs

Lentiviral constructs, LentiCas9-Blast (Addgene 52962) and lentiGuide-Puro (Addgene 52963), for CRISPR/Cas-mediated genome editing were a gift from Dr. Feng Zhang. sgRNA oligos targeting human KRAS exon 2 were designed and cloned into BsmBI site, as previously described^69^. To generate lentiviral construct for N-term GFP-tagged KRas variants, we amplified GFP-KRas WT or G12V based templates from the NCI Frederick RAS mutant collection V2.0 (Leidos). We made LV-PGK-GFP-KRAS(WT/G12V) by Gibson assembly (New England Biolabs, MA) that assembled three parts containing overlapping DNA ends into a 5.7kb lentiviral backbone.

### Protein expression and solubilization of membrane proteins in native nanodiscs from *E*.*coli*

Plasmids containing C-terminally GFP-tagged membrane proteins, SemiSWEET, AmtB, KcsA, LeuT WT and LeuT interfacial mutant (Leut-AA) were each transformed into BL21-DE3 chemically competent cells, plated onto agar plates with appropriate antibiotics, and grown overnight at 37°C. A single colony was picked from each plate and inoculated into LB media for growth with shaking at 37°C overnight. These starter cultures were used to inoculate 2 L of LB media with appropriate antibiotics. For SemiSWEET expression, the cultures were grown to an OD of 0.8, induced with 0.2 mM IPTG, and expressed overnight at 22°C (13-15 hours). For LeuT WT and LeuT-AA expression, cultures were grown to an OD of 0.3-0.4, induced with 0.1 mM IPTG, and expressed overnight at 18°C (13-16 hours). For all other proteins (AmtB, KcsA), the cultures were grown to an OD of 0.6, induced with 0.5 mM IPTG, and expressed overnight at 18°C (12 hours).

Cell pellets for SemiSWEET, AmtB, LeuT WT, and LeuT-AA were resuspended in lysis buffer (see details of all buffers used in sample preparation summarized in *Table S3*) supplemented with protease inhibitor tablets (Pierce, Thermo). KcsA expressing cell pellets were resuspended in lysis buffer supplemented with protease inhibitor tablets and PMSF (0.017 mg/ml). All cell suspensions were lysed using a microfluidizer (3 passes, 17500 psi), underwent a soft-spin to pellet cell debris following which membranes were collected via ultracentrifugation. These membranes were resuspended in 1-2% amphipathic copolymers (SMA or AASTY) in membrane resuspension buffer (details in *Table S3*) and incubated at 4°C for 2 hours for solubilization leading to formation of native nanodiscs. Solubilized membranes were then subjected to another round of ultracentrifugation to pellet insolubilized membranes. The native nanodisc-protein complexes were purified as described in later sections.

### Tissue culture

#### Culture of adherent cells

Established human PDAC cell lines (8988T, MiaPaca2, and KP-4) were sourced from DSMZ-Germany, American Type Culture Collection (ATCC), and RIKEN. All cell lines were confirmed negative for mycoplasma using PCR testing. Cells were maintained in DMEM (Corning Cellgro) supplemented with 10% fetal bovine serum (FBS) (Thermo Scientific) and 1% penicillin/streptomycin. PDAC^KRasless^ WT low and PDAC^KRasless^ G12V low cells (details on generation of PDAC^KRasless^ cells given below) were cultured using Dulbecco’s Modified Eagle Medium + GlutaMaX (DMEM, Gibco, Thermo Fisher) that was supplemented with 10% FBS (Fetal Bovine Serum, Sigma-Aldrich), Antibiotic-Antimycotic (AA, Thermo) at 100X dilution. SHSY5Y cells were cultured similarly to PDAC^KRasless^ cells, except that Advanced DMEM/F12 reduced serum medium (12634-010, Thermo) supplemented with 10% FBS and 1mM L-Glutamine was used. All cells were maintained at 37°C under 5% CO_2_.

#### Culture of suspension cells

Expi293 cells were maintained in Corning polycarbonate Erlenmeyer flasks (Thermofisher) in Expi293 expression media (Thermofisher). Cells were grown and maintained in a temperature and humidity-controlled shaker incubator at 37°C, 8% CO_2_, and 80% humidity. Sf9 cells were grown and maintained in Sf-900 III serum-free medium (SFM) (Gibco, cat. no. 10902-088) at 27°C in a humidified shaker incubator.

### Plasmid transfection/transduction

#### Transient transfection

Transient transfection was performed using standard polyethylenimine (PEI) transfection protocol. Briefly, for adherent cell lines, plasmid was diluted in OptiMEM media (Reduced Serum Media, Gibco), mixed with PEI solution diluted in OptiMEM (3 *μ*L PEI/*μ*g DNA), and incubated at room temperature for 30 minutes. This mixture was added dropwise to 10 cm tissue culture plates of 80-90% adherent cells (reagent amounts scaled for other plate sizes). For suspension cells, 1 *μ*g plasmid per mL of suspension culture was diluted into OptiMEM, while 3 uL PEI/*μ*g of DNA was separately diluted into the same volume of OptiMEM. After incubating each diluted solution for 5 minutes at RT, the diluted plasmid and PEI solutions were mixed and incubated at RT for 30 minutes. This mix was added to Expi293 cells, which were diluted to a density of 2-3 million cells/mL with fresh Expi293 expression media immediately before transfection.

#### Bacmid preparation and transduction

Bacmid was prepared in DH10Bac *E. coli* and isolated according to previously described protocol^40^. Briefly, 50 ng of plasmid was transformed into DH10Bac competent cells and plated on selection LB plates supplemented with kanamycin (50 ug/ml), Gentamycin (7 ug/ml), Tetracycline (10 ug/ml), IPTG (0.17 mM) and Bluo-gal (100ug/ml). Plates were incubated at 37 °C for ∼2 d. White colonies were used to purify bacmid DNA. To transfect Sf9 cells with bacmid, 1 × 10^6^ Sf9 cells were seeded in 6-well plates in SF-900 SFM medium. Cells were incubated at 27 °C for 20 min. For transfection, two tubes were prepared with 8 µl Cellfectin II (Thermo) in 100 µl SF-900 III SFM medium and 1 µg bacmid DNA in 100 µl SF-900 III SFM medium (Thermo). The contents of these tubes were then mixed and incubated for 30 min at RT before adding dropwise to the Sf9 cells. Cells were incubated 27 °C incubator for 72 hours. The supernatant containing the P1 virus was collected and filtered using 0.2 µm filters. The P1 virus was supplemented with 2% FBS and stored at 4°C. The P1 virus was used to generate P2 virus by inoculating 50 ml Sf9 cells, and later P2 virus was used to make P3 virus to increase the virus titer. The P3 virus titer was determined using Sf9-Easy-titer cells as described previously^40^. For TrkA expression in SH-SY5Y, cells were transduced with this P3 virus and maintained at 37 °C.

#### Lentiviral production and transduction

To produce lentivirus, we transfected plasmids that contained lentiviral backbone, packaging vector (delta8.2 or psPAX2), and envelope vector (VSV-G) into HEK293T cells using TransIT-LT1 (Mirus Bio). Supernatant post-48/72h transfection was collected for transduction in target cells supplemented with 8 µg/mL polybrene (EMD Millipore). Single cell clones of Cas9 and sgKRAS-transduced cells were sorted into 96-well plates using a FACSMelody benchtop sorter (BD Biosciences) and verified for successful knock-out of KRas by Western blot analysis and PCR amplicon sequencing. To acquire cell populations with low or high levels of GFP-Kras expression, cells were sorted based on the fluorescence intensity of GFP by FACSMelody (BD Biosciences). Fusion protein expression was further validated by Western blotting.

### Cellular treatments with small molecules and growth factor

For relevant KRas oligomerization studies (Figure 4), cells were treated with either 10 µM BI-2852 or an equivalent volume of solvent (DMSO) for 2 hours immediately before harvesting. 10 µM BI-2852 was maintained throughout the copolymer solubilization and purification process. In relevant TrkA oligomerization studies, SH-SY5Y cells were serum starved for 16 hours, and then either harvested as is or after treatment with NGF at a concentration of 100 ng/ml for 15 min. For these studies, a TrkA specific kinase inhibitor (GW441756, IC_50_ = 2 nM)^70^ was maintained at a concentration of 5 µM throughout expression.

### Protein expression and solubilization of membrane proteins in native nanodiscs from mammalian cells

#### Solubilization of membrane proteins from Expi293 expression cultures

KRas was expressed in Expi293 for 16 hours before harvesting the cells. Cells were resuspended in lysis buffer (*Table S3*) supplemented with protease inhibitor cocktail tablets (Pierce, Thermo) and lysed using nitrogen cavitation (600 psi, 20 minutes). Debris and cell nuclei were removed by a soft spin and clarified lysate was ultracentrifuged to isolate the membrane fraction. Membranes were solubilized using a 1-2% amphipathic copolymer (SMA or AASTY) in membrane resuspension buffer (*Table S3*) at 4°C. Solubilized membranes were then subjected to another round of ultracentrifugation to pellet any undissolved membrane, isolating soluble native nanodiscs in the supernatant.

#### Solubilization of membrane proteins from adherent cell cultures

PDAC^KRasless^ cells stably expressing KRas WT or G12V were seeded into 15-cm plates to confluency in 24 hours, and cells were harvested. For TrkA expression, SH-SY5Y cells transduced with P3 baculovirus were harvested from 10 cm plates (one plate per experiment condition) after about 24 hours of expression. Cells were lysed by resuspending in lysis buffer and using a Dounce homogenizer (20-30 strokes). Further processing of samples to generate native nanodiscs was identical to the procedure outlined above.

### Native nanodisc purification

#### Affinity chromatography

Native nanodiscs containing KRas were purified using affinity chromatography (Ni-NTA resin, Cube biotech) at 4°C. Samples were adjusted with imidazole for Ni-binding, washed extensively with wash buffer, and eluted with elution buffer using gravity flow columns. Buffer compositions were same as membrane resuspension buffer, except they were supplemented with 5 mM imidazole (binding buffer), 15 mM imidazole (wash buffer), and 350 mM imidazole (elution buffer). Purified nanodisc samples were subjected to FSEC as described below.

#### Fluorescence size-exclusion chromatography

Each native nanodisc sample was subjected to fluorescence detection size-exclusion chromatography (FSEC) on a Superose 6 10/300 GL column (Cytiva) either immediately after solubilization (SemiSWEET, AmtB, KcsA, LeuT variants, and TrkA) or following affinity purification (for KRas) using an AKTA Pure FLPC (GE). A Shimadzu RF-20A XS fluorescence detector was installed on-line to the FPLC, allowing detection of GFP (488 nm) signal. The recipes of the FSEC buffer for each protein are detailed in *Table S3*. Non-void volume fractions containing the GFP-positive native nanodisc target protein complex were collected and subjected to Native-nanoBleach analysis.

### Single-molecule imaging

#### Device assembly and glass substrate functionalization

All single-molecule experiments were performed in flow chambers (sticky-Slide VI 0.4, Ibidi, Planegg, Germany) that were affixed to functionalized glass substrates. Glass substrates were generated by first cleaning glass slides (Ibidi glass coverslips, bottom thickness 170 µm+/–5 µm) by bath sonication in 2% Hellmanex III solution (Hellma Analytics) for 30 min at 37°C. This was followed by extensive washing with ddH_2_O, and bath sonication for 30 min in a 1:1 mixture (vol/vol) of isopropanol:water. The glass slides were washed again with ddH_2_O, then air dried and cleaned for 5 min in a plasma cleaner (Harrick Plasma PDC-32 G, Ithaca, NY). Immediately after plasma cleaning, flow chambers were affixed to these glass slides to assemble the device.

After assembly, the glass substrates were treated with a mixture of Poly-L-lysine PEG and PEG-Biotin (PEG-biotin diluted 1:200 in PLL-PEG, stock concentrations 1 mg/mL) for 30 min (SuSoS, Dübendorf, Switzerland). The glass substrates were then washed with 3 mL of Dulbecco’s phosphate-buffered saline (DPBS, Gibco, Thermo Fisher). Streptavidin (1 mg/mL, NEB N7021S) was diluted 1:10 in DPBS and applied to passivated flow chambers for 30 min. Following incubation, excess streptavidin was washed 3 mL with DPBS. Next biotinylated GFP-nanobody (1:100 dilution in dilution buffer - 25 mM Tris pH 8.0, 150 mM KCl, 1 mM TCEP; stock concentration 1 mg/mL, ChromoTek gtb250) were applied to the flow chambers, which had been equilibrated with 3 mL dilution buffer, and incubated for 30 min. Subsequently, excess nanobody was washed with 3 mL dilution buffer and then 3 mL DPBS. All incubations were done at room temperature (RT).

#### Sample capture at single molecule density

Prior to immobilization, native nanodisc samples were diluted in the corresponding FSEC buffer such that the capture on to the glass substrate results in optimal single-molecule density. To immobilize native nanodiscs for imaging, 200 *μ*L of each native nanodisc sample was applied to the functionalized substrate and incubated for 5 minutes at RT and then washed with 3 mL DPBS.

#### Image acquisition

Single-particle TIRF images were acquired on a Nikon Eclipse Ti-inverted microscope equipped with a Nikon 100X 1.49 numerical aperture oil-immersion TIRF objective, a TIRF illuminator, a Perfect Focus system, and a motorized stage. Images were recorded using an Andor iXon electron-multiplying charge-coupled device camera. The sample was illuminated using the LU-N4 laser unit (Nikon, Tokyo, Japan) with solid state lasers for the 488 nm, 561 nm and 640 nm channels. Lasers were controlled using a built-in acousto-optic tunable filter (AOTF). The 405/488/561/638 nm Quad TIRF filter set (Chroma Technology Corp., Rockingham, Vermont) was used along with supplementary emission filters of 525/50 m, 600/50 m, 700/75 m for 488 nm, 561 nm, 640 nm channel, respectively.

Samples were imaged using 3-color acquisition to assess the density of molecules within the imaging area, and to confirm the lack of background signal in 561 nm and 640 nm channels for samples containing only GFP-tagged protein. This was performed by computer-controlled change of illumination and filter sets (488 nm, 561 nm, and 640 nm) at 20 different positions from an initial reference frame, so as to capture multiple non-overlapping images. Native nanodiscs were imaged by setting the 488 nm laser to 20 mW, 561 nm laser to 56 mW, and 640 nm laser to 20 mW, respectively using an exposure time of 80 milliseconds for all three channels. For photobleaching experiments, the field of view was illuminated in the TIRF mode by laser at 488 nm (GFP fluorescence) and signal was recorded in a stream acquisition mode to collect movies over 1500 frames at ∼9.3 frames/sec, with an exposure time of 80 milliseconds and with the 488 nm laser power set to 1.8 mW and 2.7 mW for each sample. All mW values refer to power at the laser source. All image acquisition was carried out using the Nikon NIS-Elements software.

#### Image analysis for GFP-photobleaching step counting

Individual single particles were detected and localized using the single particle tracking plugin TrackMate in ImageJ^71–73^. The particles were localized with the Difference of Gaussian (DoG) detector with an initial diameter set to six pixels, and the detection threshold value set to 50. Particles outside the center area of 350 × 350 pixel^2^ were excluded due to heterogeneous TIRF illumination. No further filtering processes were applied in TrackMate.

Frames from the GFP step-photobleaching movies were analyzed to plot the time-dependent decrease in GFP fluorescence intensities for each tracked particle in the field of view using a custom MATLAB code^74^. These plots were analyzed manually by scoring them to have 1-4 photobleaching steps or discarded if no clear photobleaching steps were identified. Photobleaching counts corresponding to 1000-1500 individual particles were recorded for each biological replicate, and reported as mean +/-sd over three to four biological replicates for each sample. The step counting was also verified through average GFP intensity for each particle. This step distribution data over 1-4 steps for each sample was converted into a corresponding distribution of monomers, dimers, trimers, and tetramers, considering the GFP maturation efficiency to be 70%, as described below.

### Calculation to convert GFP photobleaching step distribution to oligomeric distribution

The number of photobleaching steps underrepresents the number of subunits within an oligomer due to the ∼70% maturation efficiency of GFP^11,41^. For example, 1-step photobleaching may arise from monomers but also from some dimers with only one subunit carrying a mature, fluorescent GFP. We accounted for the dark GFP state as follows. We first obtained the theoretical probability distribution describing how many subunits are GFP fluorescent within a given oligomeric state (from monomer to tetramer).

Let us represent the GFP maturation probability as *p* = 0.7.

The theoretical probability of an oligomer with *n* subunits bleaching with exactly *k* steps:

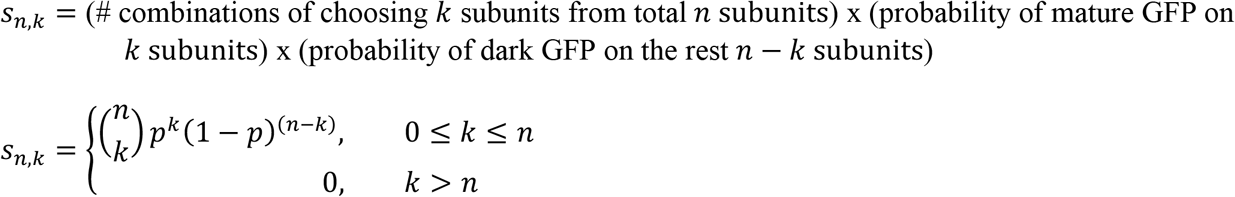

For each oligomer with n subunits (*n* = 1 - 4), using the above equation, we obtained a theoretical distribution of 0, 1, 2, …, *n* steps. In any single-molecule imaging technique, we only experimentally observe a signal if at least one of the subunits carry a mature GFP within the oligomer. Hence, we remove the percentage of 0-steps from this theoretical step distribution and normalize the fractions of 1, 2, …, *n* steps such that they represent 100% of the experimentally observed spots. Therefore, the theoretical step distribution for an oligomer is given by the equation:

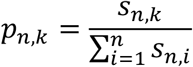

The normalized theoretical fractions of *k* steps (i.e., theoretical step distribution) for monomers, dimers, trimers, and tetramers are tabulated below.

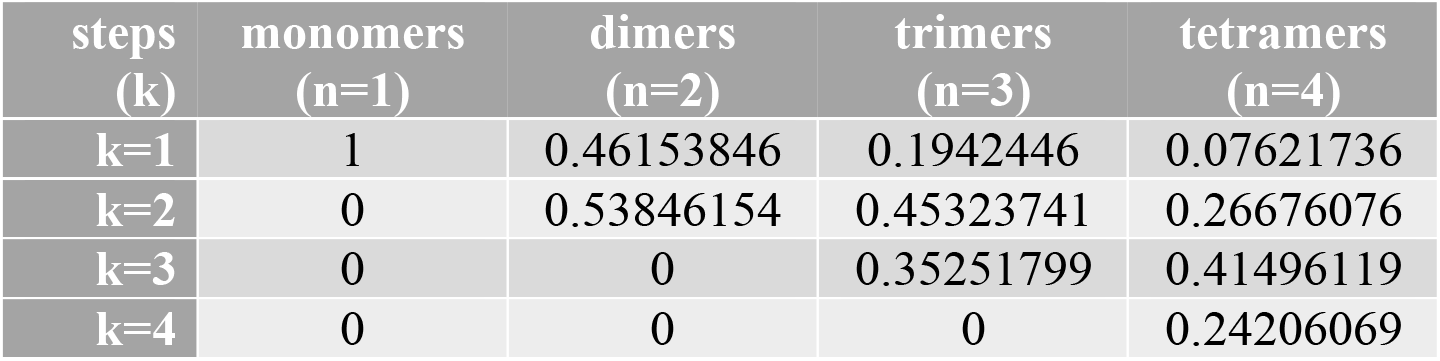

| steps<br>(k) | monomers<br>(n=1) | dimers<br>(n=2) | trimers<br>(n=3) | tetramers<br>(n=4) |
| --- | --- | --- | --- | --- |
| k=1 | 1 | 0.46153846 | 0.1942446 | 0.07621736 |
| k=2 | 0 | 0.53846154 | 0.45323741 | 0.26676076 |
| k=3 | 0 | 0 | 0.35251799 | 0.41496119 |
| k=4 | 0 | 0 | 0 | 0.24206069 |

For proteins exhibiting a single stoichiometry (i.e., either monomer, dimer, trimer, or tetramer), we can directly match the experimental step distribution with one of these theoretical step distributions to identify the specific oligomeric state (*Fig. 2a*). However, for proteins with mixed oligomeric distributions, we further calculated the relative fractions of monomers, dimers, trimers, and tetramers whose combined theoretical distribution best fit the experimentally-obtained step distribution. The calculation for this conversion is given below.

Let us represent the experimentally-obtained fraction of *k* bleaching steps as *o*_*k*_ and the unknown fractions of monomer, dimer, trimer, tetramer as *x*_1,_ *x*_2,_ *x*_3,_ *x*_4_, respectively. Then the system of 4 linear equations that explain the experimentally obtained fractions *o*_*k*_ is as follows:

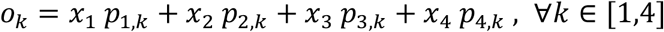

Considering that *x*_1_ + *x*_2_ + *x*_3_ + *x*_4_ = 1, this equation can be simplified to 3 linear equations with 3 unknown variables as below:

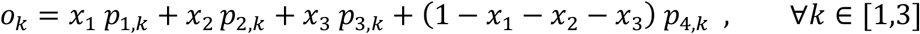

This can be rearranged to the following:

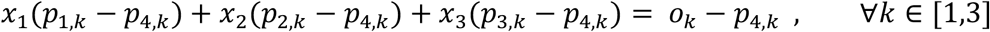

These equations with three unknown variables (*x*_1,_ *x*_2,_ *x*_3_) can be represented as matrix equation *AX* = *O* − *C* where:

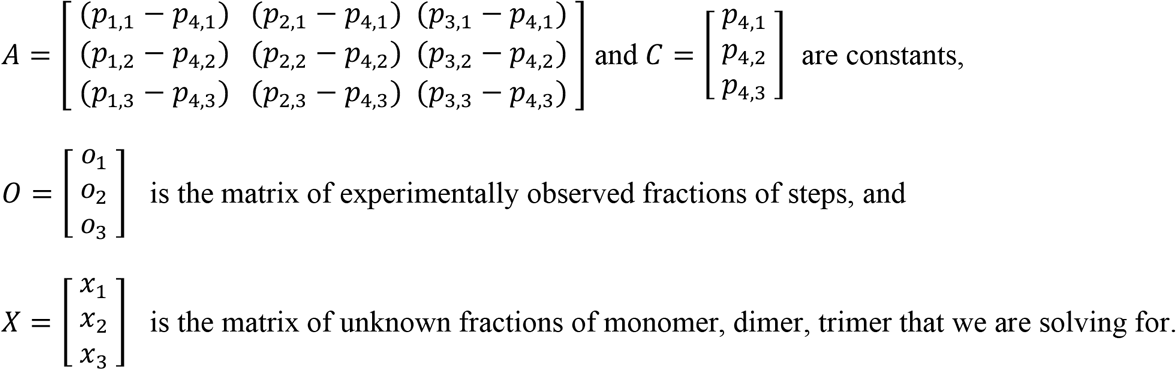

Hence, given an observed values for *O*, we can solve for the fraction of monomers, dimers, and trimers as

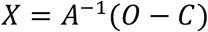

The fraction of tetramer is then calculated as *x*_4_ = 1 − (*x*_1_ + *x*_2_ + *x*_3_). We wrote a Matlab code to do this calculation.

### Confocal imaging

PDAC^KRas-GFP^ (low) WT/G12V cells were plated at 20,000 cells/mL into 35 mm Mattek dishes (P35GC-1.5-14-C) and incubated overnight. After 24 hour the cells were exchanged into DPBS and imaged with a Dragonfly 505 spinning disc confocal (Andor) mounted into a Ti2-E inverted microscope (Nikon) with a plan apochromat lambda 60X oil, NA 1.42 objective and a SONA 4.2MP camera (SONA-4BV6X; 102nm/pixel; Andor) under the control of Fusion (2.4.0.6, Andor). Images were acquired using 50% 488 nm laser power with 155 milliseconds exposure, and with 2×2 binning.

### Co-immunoprecipitation

Expi293 cells expressing combinations of GFP-KRas and/or FLAG-KRas, treated with either DMSO or 10 *μ*M BI-2852, were harvested and flash frozen. Cells were then lysed and solubilized with copolymers as described above. Native nanodisc samples were subjected to immunoprecipitation using anti-FLAG M2 resin (M2 anti-FLAG affinity resin, Sigma) according to manufacturer’s instructions. Native nanodisc samples were incubated with equilibrated anti-FLAG resin for 2 hours at 4°C. Beads were pelleted by centrifugation and the flow through (supernatant) was collected. The beads were washed 3 × 1 mL with membrane resuspension buffer for KRas before the beads were eluted with membrane resuspension buffer supplemented with 0.2 mg/mL 3X FLAG peptide (SAE0194, Sigma). Aliquots of samples were collected at each stage (input to anti-FLAG resin, flow through, and eluate) and analyzed using western blotting as described below.

### Western blot

#### General WB sample preparation

Cells were harvested and resuspended in lysis buffer supplemented with phosphatase inhibitor cocktail (*Table S3*, for TrkA the lysis buffer was also supplemented with 1% DM) and lysed using a Dounce homogenizer. Lysates were spun at 1600 x g for 8 minutes on a table top centrifuge to remove large debris/nuclei. For western blotting analysis, samples were diluted in 4X laemmli buffer (supplemented with 2-mercaptoethanol) and denatured at 95°C for 5 minutes. For in-gel GFP fluorescence analysis, samples were prepared as outlined above, except no reducing agent (2-mercaptoethanol) was added to laemmli buffer, and the samples were not heat denatured before loading onto polyacrylamide gels.

For comparison of KRas levels (see Figure 5), cells were lysed with ice-cold RIPA buffer (Pierce), supplemented with 0.5 µM EDTA and Halt protease and phosphatase inhibitors (Thermo Scientific). Lysates were rotated at 4°C for 15 min to mix, and centrifuged at maximum speed for 10 min to collect whole cell lysates. Protein concentration was measured with the BCA protein assay (Pierce), and samples were diluted in 4X laemmli buffer supplemented with 2-mercaptoethanol.

#### Western blot

For Western blotting, samples were loaded onto 12% Tris-glycine gels (#1658015, Biorad). Proteins were transferred onto ethanol-activated PVDF membranes (#1620177, Biorad). The following antibodies were used for immunoblotting: rabbit anti-Na/K ATPase (Abcam ab167390, 1:5000), rabbit anti-calnexin (Abcam ab92573, 1:2000), rabbit anti-RAS (Abcam ab52939, 1:1000), rat anti-FLAG (Sigma SAB4200071, 1:1000), rabbit anti-GFP (CST 2956, 1:1000), rabbit anti-TrkA (CST 2505S, 1:2000), rabbit anti-TrkA-pY490 (CST C35G9, 1:2000). Primary antibodies were detected with species-specific secondary antibodies linked to HRP for chemiluminescence detection (see below). Na/K ATPase and calnexin were used as loading controls. Blot were developed using SuperSignal West Dura (Thermo Scientific, 34076). For comparing KRas levels in PDAC and PDAC^KRasless^ cell lines (see Figure 5), an aliquot of 25µg of total protein per sample was loaded into 4–20% Mini-PROTEAN® TGX™ precast gels (Bio-Rad) for SDS-PAGE separation. Proteins were transferred to nitrocellulose membranes using the Trans-Blot® Turbo™ transfer system (Bio-Rad). The following antibodies were used for immunoblotting: rabbit anti-HSP90 (CST 4877, 1:1000), rabbit anti-GFP (CST 2956, 1:1000), mouse anti-KRAS (SCBT sc-30, 1:200), rabbit anti-pERK1/2(T202/Y204) (CST 4370, 1:1000), mouse anti-ERK1/2 (CST 9107, 1:1000), rabbit anti-pAKT(S473) (CST 4060, 1:2000), mouse anti-AKT (CST 2966, 1:2000). Mouse anti-HSP90 (BD 610418, 1:10,000) was used as loading control.

For in-gel GFP fluorescence, after separating proteins using SDS-PAGE, the gels were washed 2X with about 50 mL of ddH_2_O before in-gel fluorescence was imaged (see below).

#### Western blot image acquisition and analysis

To image Western blots, images of target bands were acquired using either the chemiluminescence or fluorescence channels (based on secondary antibody used) on a Biorad ChemiDoc MP system. Protein standards were imaged using the colorimetric channel, and these images were merged with target band images to detect molecular weights of observed bands. Western blots were quantified using the Gel Analyzer function in Fiji image analysis software^73^. Briefly, bands were selected using the rectangular selection tool, line profile peaks were generated and delineated using the line tool, then each peak area was quantified using the wand tool.

In-gel fluorescence was imaged by laying gels directly on the appropriate Biorad imaging tray and acquiring images in the alexa-488 channel. Protein standards were imaged using the Coomassie blue channel, and these images were merged with alexa-488 fluorescence images to allow for visualization (due to the non-denaturation of proteins within these gels, proteins sometimes migrated differently than would be expected given their molecular weights). Gels were analyzed using the Gel Analyzer function of Fiji, as described above. All gel or blot images are shown at the same scale, and all quantified images were acquired using same exposure times.

### Negative-stain electron microscopy

Negative staining was performed according to the method outlined in^75^. Briefly, glow-discharged grids (Carbon Type-B, 400 mesh, Cu, Ted Pella) were floated on a 5 µL sample bead on parafilm for 1 minute. Grids were washed by floating on 4 consecutive beads of 0.75% (w/v) uranyl formate (UFO), immediately wicking residual liquid with filter paper before placing the grid on each drop. Samples were stained by floating the grid on a droplet of UFO for 20 s. After removing all stain by wicking thoroughly with a filter paper, the grids were air-dried and stored in a grid box.

Images were acquired on a Tecnai T12 microscope (FEI) equipped with a LaB6 filament and operated at an acceleration voltage of 120 kV. Images were recorded using a Gatan Ultrascan 4000 (4k x 4k) CCD camera at 52,000X magnification and a defocus of −1.5 µm to -5 µm. The pixel size in resulting images was 2.14 Å/pixel. To quantify the size distribution of native nanodiscs, images were analyzed in Fiji by using the line tool to draw straight lines extending the length of the diameter of each individual native nanodisc that was present in multiple EM images (images over different areas of the grids were used to produce at least 1000 measured diameters). Each of these lines was added to the ROI manager, and the length of each line (nm) stored in the ROI manager was tabulated using the measure function in Fiji. The native nanodiscs diameters were visualized as a histogram (bin size = 1 nm), and the data was fit to a gaussian distribution using the Prism 9 software.

### Theoretical calculation for correlating probability of chance overlap as a function of membrane protein surface density

We calculated the theoretical probability of chance overlap between two or more protein subunits on a 10 nm X 10 nm area (for spatial resolution of ∼10 nm) over a wide range of protein surface densities (in molecules/*μ*m^2^) as described below.

Let us assume the surface density of a protein is *m* molecules per *μ*m^2^. We divide the 1 *μ*m^2^ area into 10 nm X 10 nm bins and calculate the probability of *k* molecules (where *k* = 0, 1, 2,…) falling in a given bin.

The number of 10 nm x 10 nm bins within 1 *μ*m^2^ (i.e., 1000 nm X 1000 nm) area: *b* = 10^6^ nm^2^ / 100 nm^2^ = 10000 The probability of a molecule falling in a given bin: *p* = ^1^/*b*

The probability of exactly *k* molecules falling in a given bin:

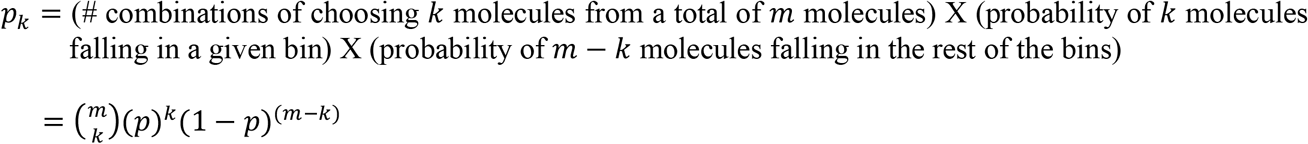

The probability of two or more molecules falling in a given bin, i.e., the probability of chance overlap within a given area on the membrane surface: *p*_2c_ = 1 − *p*_d_ − *p*_1_, where *p*_d_ and *p*_1_ are the probabilities that zero or only one molecule will fall within a defined bin.

In other words, the probability of any 10 nm x 10 nm area to have overlapping molecules when the surface density is *m* molecules per um^2^ is:

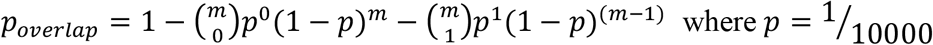

We wrote a Matlab code to do this calculation over a wide range of membrane protein surface densities (in molecules/*μ*m^2^) and for four different spatial resolutions (10, 20, 30, and 200 nm), which in our calculation determines the bin size (10 nm X 10 nm in the calculation above) and, therefore, the number of bins within a 1 *μ*m^2^ area. We used GraphPad Prism to plot the traces.

## Data availability

All single-molecule imaging data generated and analyzed during the current study are available from the corresponding author upon request.

## Code availability

The calculations detailed in the Methods section above have been formulated as Matlab codes that are available from the corresponding author upon request.

## Acknowledgements

We thank members of the Bhattacharyya, Gupta, and Muzumdar labs for helpful discussions. We especially thank Anthony Quinnert (Bhattacharyya lab) for maintenance of our microscopy setup and Felix Rivera-Molina for help with confocal microscopy. We thank Marc Llaguno for help with negative stain EM data collection. We also thank members of the Pharmacology department at Yale University for helpful discussions on this work and feedback on our manuscript. The pQE60-KcsA construct was a gift from Dr. Crina Nimigean’s lab. The pET16b-LeuT construct was a gift from Dr. Eric Gouaux’s lab. Expi293 cells were a gift from Dr. Karin Reinisch’s lab. GW acknowledges support from the PPTP training grant (T32-GM007324), CB acknowledges support from the NSF GRFP fellowship (DGE-2139841), and SK acknowledges support from NIH grants (R00GM126145 and R35GM147095). X.G. was a CSC-Yale Scholar. M.D.M. acknowledges support from an NCI Mentored Clinical Scientist Research Career Development Award (K08-CA208016), a NIH New Innovator Award (DP2-CA248136), a Lustgarten Foundation Therapeutics Focused Research Program award, an American Cancer Society Institutional Research Grant (#IRG 17-172-57), and in part, the Yale Comprehensive Cancer Center Support Grant (P30CA016359). KG acknowledges support from NIGMS (R01GM141192). MB acknowledges support from NIGMS (R00GM126145 and R35GM147095) for funding.

