## Supplementary Figures and Tables for "Determination of oligomeric organization of membrane proteins from native membranes at nanoscale-spatial and single-molecule resolution"

**Supplementary Figures and legends**

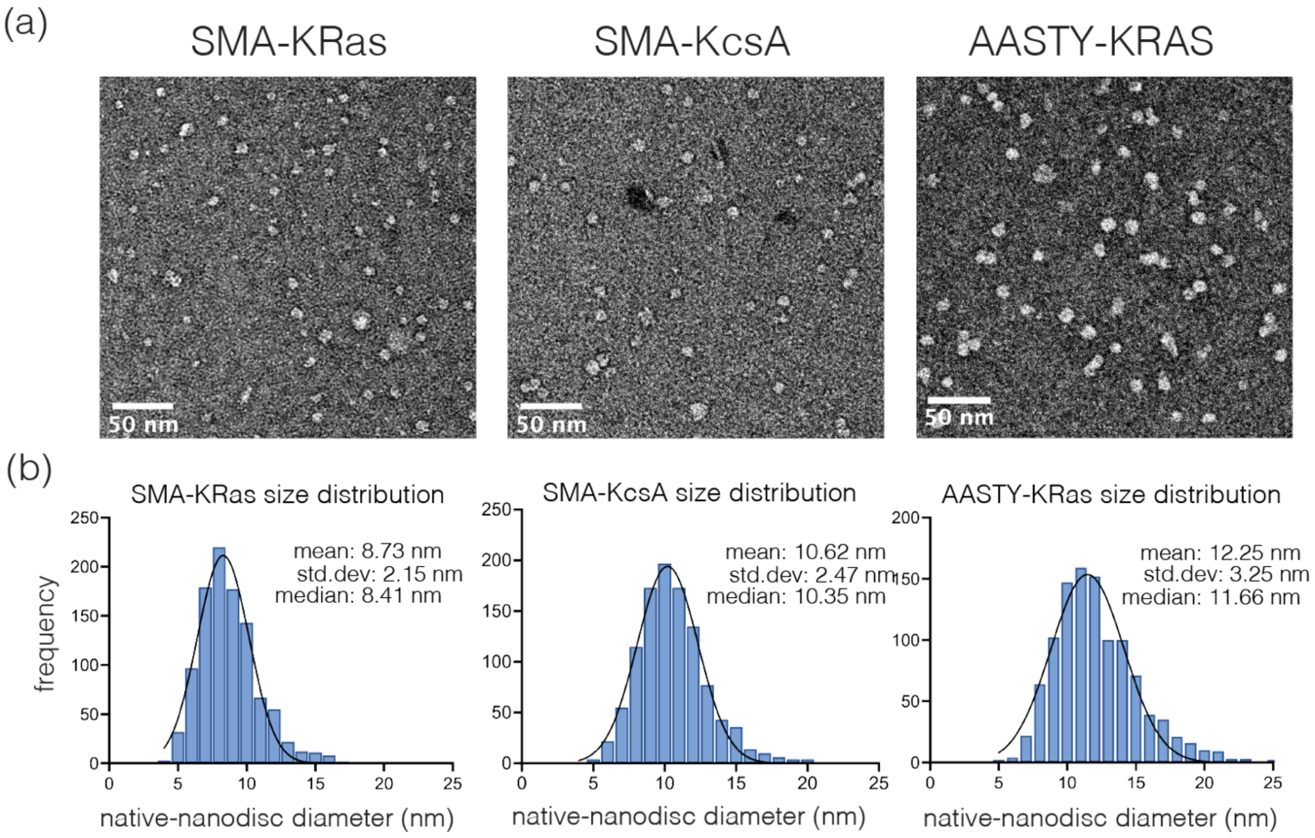

**Supplementary Figure 1:** Size distribution of native nanodiscs using negative-stain EM. (a) Representative negative stain electron microscopy images depicting native nanodiscs encapsulating GFP-tagged KRAS and KcsA obtained using SMA or AASTY. (b) Plot showing the distribution of native nanodisc-encapsulated KRas and KcsA diameters (in nm) measured over ~1000 individual particles.

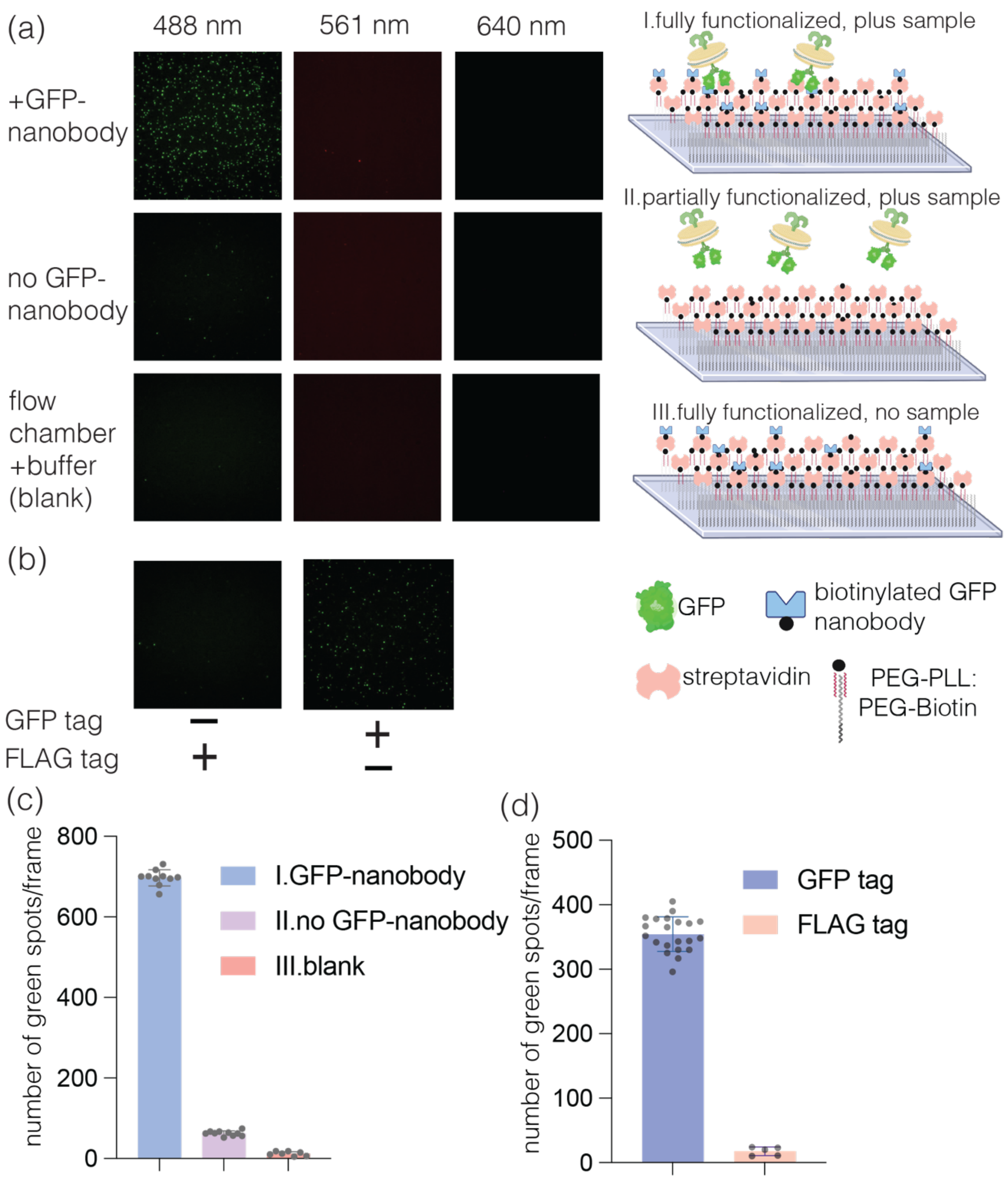

**Supplementary Figure 2:** Immobilization of native nanodiscs on glass substrates at single-molecule density. (a) Representative single-molecule TIRF images showing immobilization of native nanodiscs. Flowing GFP-KRas native nanodisc samples over a fully functionalized substrate leads to efficient nanodisc capture (+ GFP-nanobody, top panel). Flowing the same samples over a partially functionalized substrate that has been treated up to the streptavidin step but not coated with biotinylated GFP-nanobody (middle panel) shows almost no signal when excited by the 488 nm wavelength. The 561 nm and 640 nm channels yield no signal under our experimental conditions, showing a clean background. A control with a fully functionalized substrate but no sample (blank well, bottom panel) with buffer alone shows a clean background in all three fluorescent channels (488, 561, 640 nm). Cartoon depiction of the experimental conditions are shown on the right of each panel. (b) Representative single-molecule TIRF images showing immobilization of native nanodiscs by flowing GFP-KRas or FLAG-KRas (dark) native nanodisc samples over fully functionalized substrates. Capture was seen only for GFP-KRas containing samples. All images in (a) and (b) are scaled equally. (c) Number of fluorescent spots in the 488 nm channel (GFP) per imaging frame for data shown in (a). (d) Number of fluorescent spots in the 488 nm channel (GFP) per frame for data shown in (b). Data in (c-d) are presented as mean +/- sd for 7-20 frames.

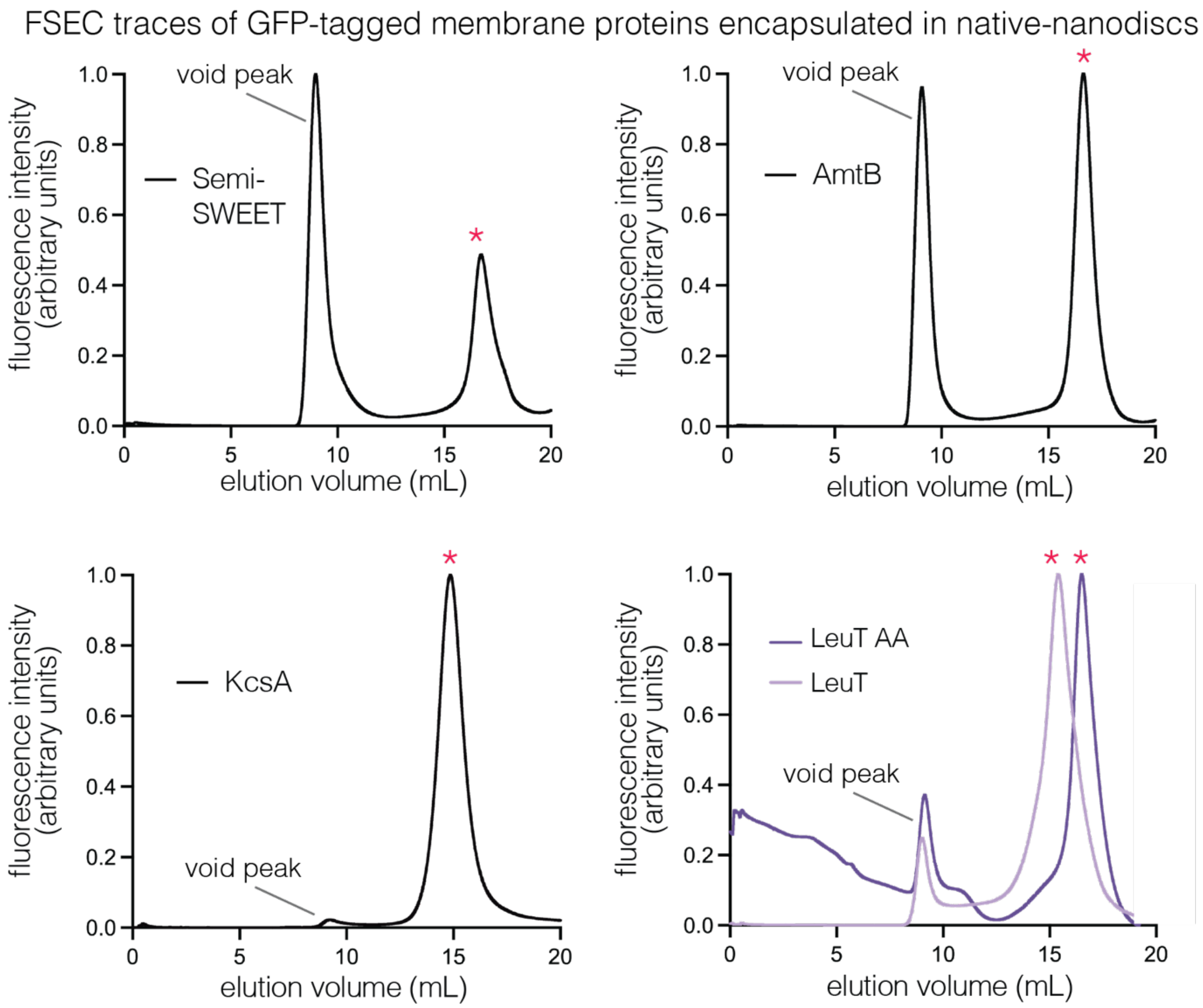

**Supplementary Figure 3:** FSEC profiles of native nanodiscs encapsulating diverse membrane proteins with well-established oligomeric states – semiSWEET (dimer), AmtB (trimer), KcsA (tetramer). Overlay plot depicts FSEC traces of native nanodiscs containing LeuT or LeuT AA, a mutant at residues F488 and Y489 at the LeuT dimeric interface. GFP fluorescence is monitored and the non-void volume, GFP-positive fractions were collected for analysis (marked by a red *). The FSEC traces are normalized to have the same area under the curve.

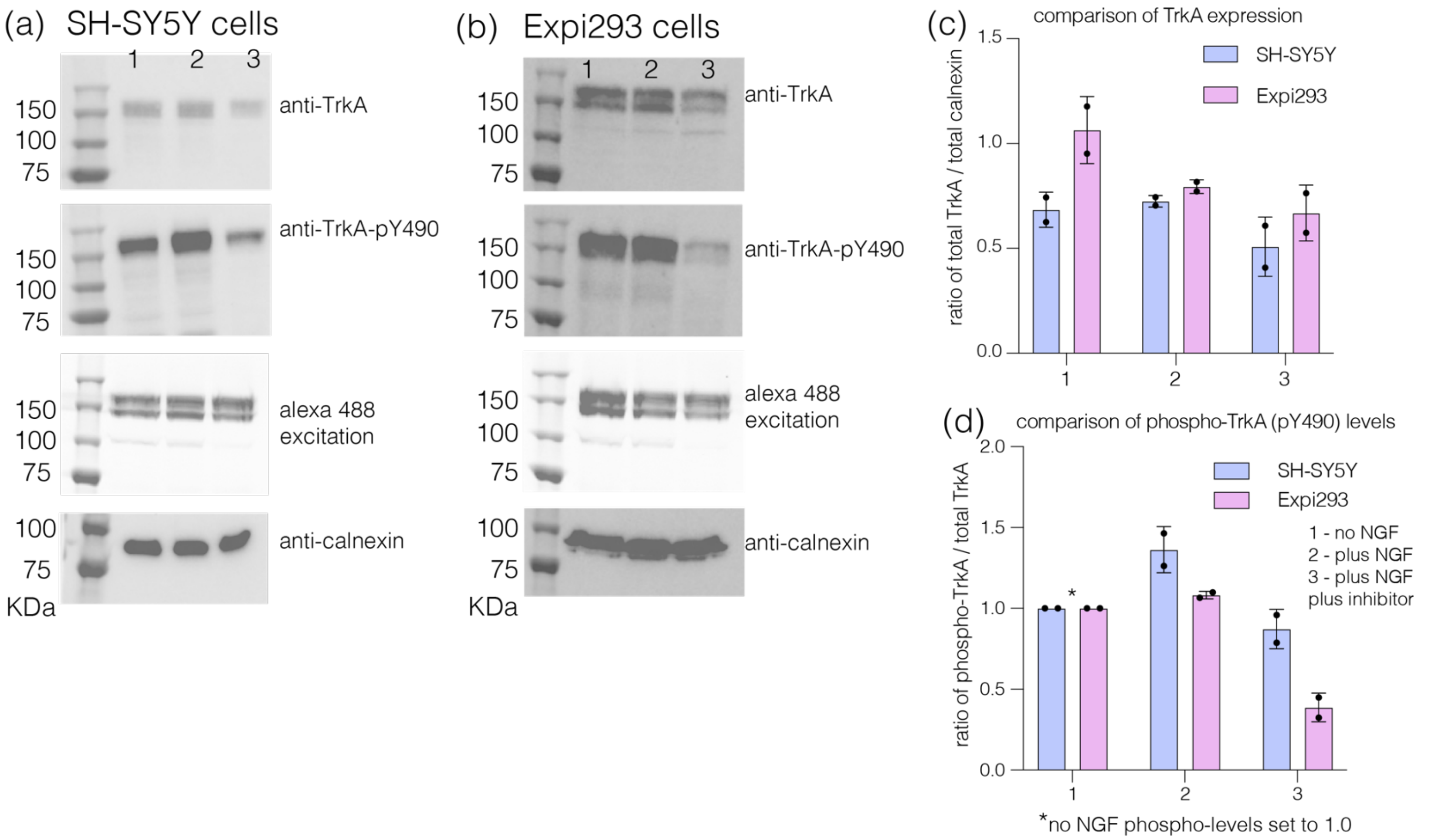

**Supplementary Figure 4:** Expression and autophosphorylation of TrkA in Expi293 and SH-SY5Y cells. (a) Representative Western blots showing the expression levels of TrkA (using anti-TrkA antibody CST 2505S as well as SDS gel excited with 488 nm) and the extent of TrkA autophosphorylation at Y490 (phospho-specific Y490 antibody CST C35G9) in SH-SY5Y cells expressing GFP-tagged TrkA. The total amount of calnexin in the membrane was used as a loading control (Abcam ab92573). Data is shown for cells under three different conditions: 1 – no NGF, 2 – cells treated with 100 ng/mL NGF, and 3 – cells treated with 100 ng/mL NGF and 5 μM of TrkA-kinase inhibitor. (b) Same as (a) for Expi293 cells expressing GFP-tagged TrkA. (c) Comparison of the amount of GFP-TrkA expressed in SH-SY5Y and Expi293 cells under conditions 1-3. TrkA expression levels in each cell line were quantified using band densitometry and normalized with respect to total amounts of calnexin using Fiji. (d) Comparison of the total phospho-TrkA levels (pY490) in SH-SY5Y and Expi293 cells under conditions 1-3. Autophosphorylation levels at Y490 in each cell line were quantified using band densitometry and normalized with respect to total amounts of TrkA expressed in the corresponding cell lines using Fiji. All primary antibodies were used at 1:2000 dilution, the exposure for acquiring the western blot images was 15 sec, and the 488 nm excitation was 5 sec for all samples. Data is shown as mean ± sd.

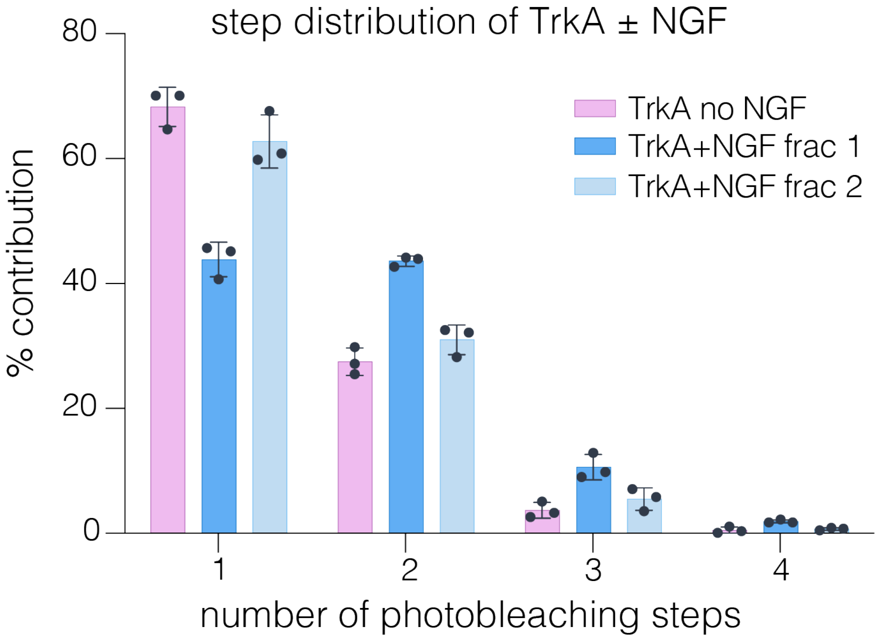

**Supplementary Figure 5:** Experimentally obtained GFP-photobleaching step distribution of TrkA in the presence and absence of NGF. Data is shown from a total of about 3000-4000 spots over at least three biological replicates as mean ± sd.

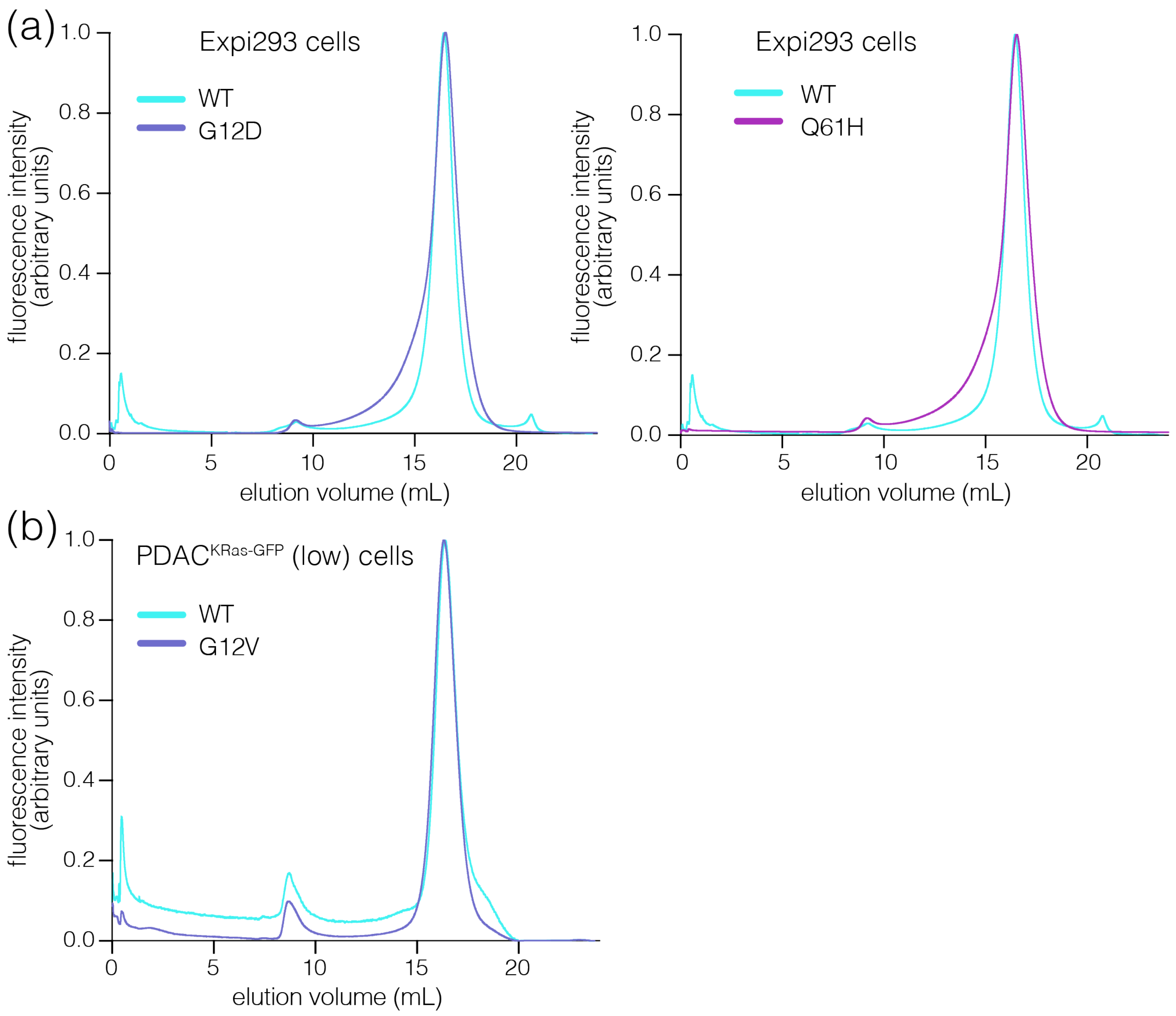

**Supplementary Figure 6:** FSEC profiles of KRas variants (WT and oncogenic mutants) from Expi293 and PDAC cells. (a) Overlay of FSEC traces from native nanodiscs containing KRAS WT and its oncogenic mutants G12D or Q61H isolated from Expi293 cells. (b) Overlay of FSEC traces from native nanodiscs containing KRAS WT and G12V isolated from PDAC$^{KRAS\text{-}GFP}$ (low) cells. The plots show GFP fluorescence intensity when excited at 488 nm. The FSEC traces are normalized to have the same area under the curve.

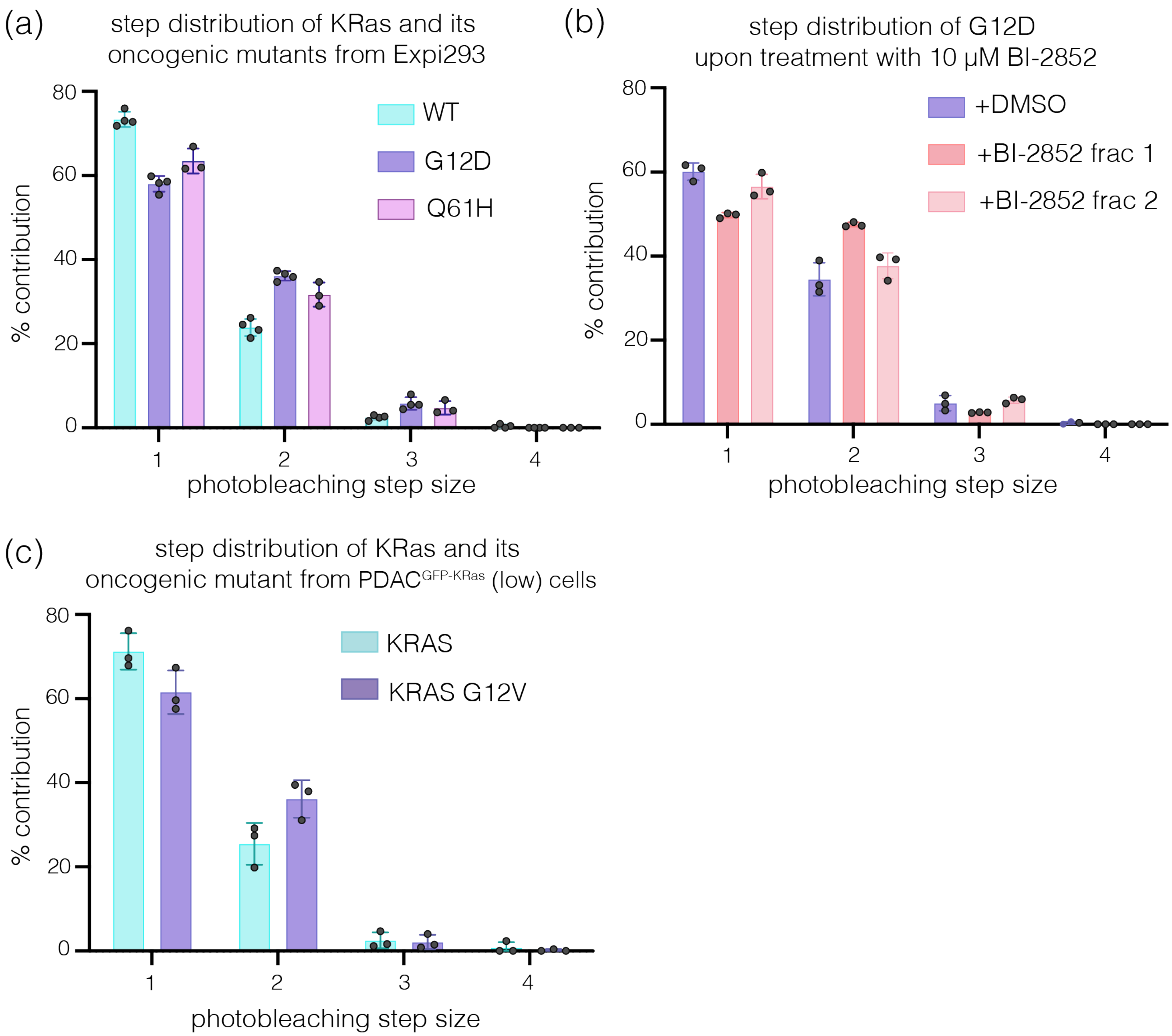

**Supplementary Figure 7**: Photobleaching step distributions obtained from KRas containing native nanodiscs. (a, c) Experimentally obtained GFP-photobleaching step distribution of KRas WT and its oncogenic mutants in native nanodiscs isolated from Expi293 and PDAC$^{KRAS\text{-}GFP}$ (low) cells, respectively. (b) Experimentally obtained GFP-photobleaching step distribution of KRas G12D in native nanodiscs isolated from Expi293 cells that were treated with DMSO alone (control) or 10 μM BI-2852 solution in DMSO. All photobleaching step-analyses shown in (a-c) are from a total of about 3000-4000 spots over at least three biological replicates. Data shown as mean ± sd.

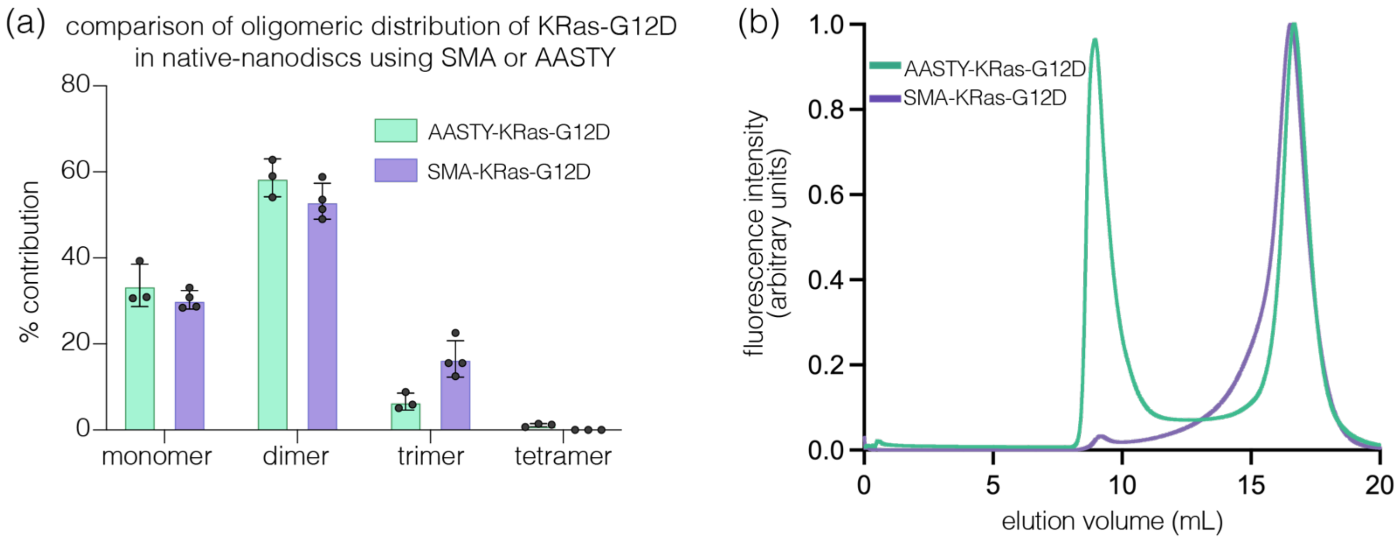

**Supplementary Figure 8**: Native-nanoBleach analysis of KRas in SMA or AASTY nanodiscs. (a) Oligomeric distribution of GFP-KRas oncogenic mutant G12D within native nanodiscs isolated from Expi293 cells using two different amphipathic co-polymers – SMA and AASTY. Data shown for a total of about 1200-4000 spots over 2-4 biological replicates and shown as mean ± sd. (b) FSEC traces for native nanodisc-KRas obtained using SMA and AASTY. The FSEC traces are normalized to have the same area under the curve.

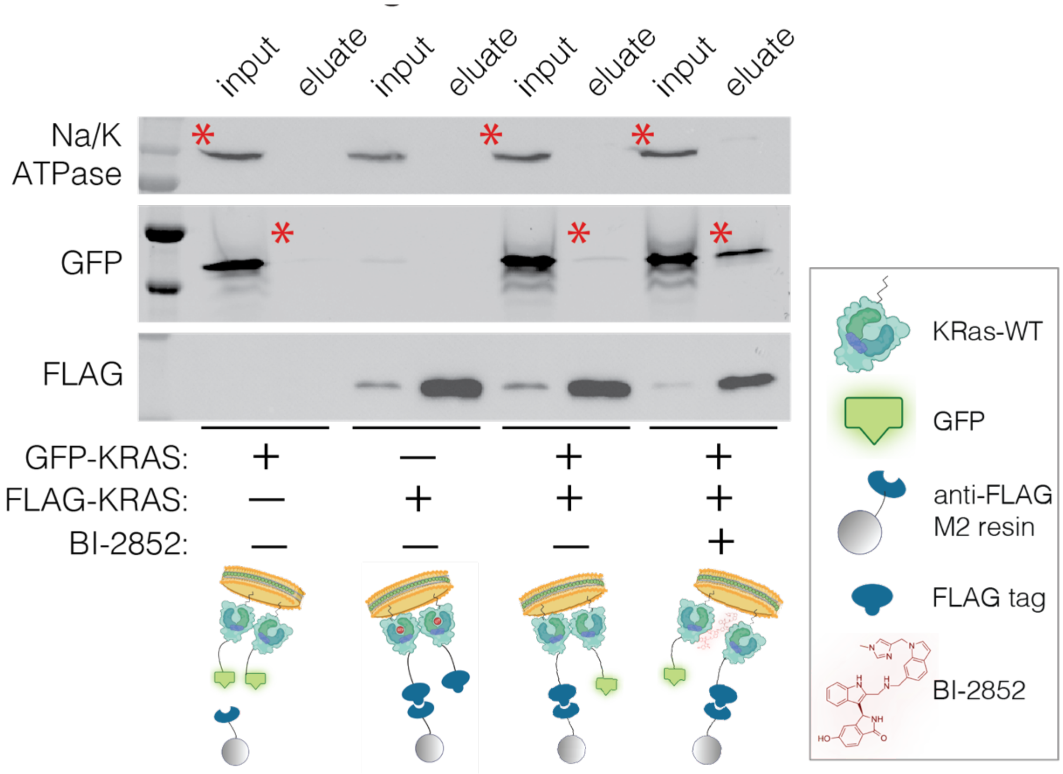

**Supplementary Figure 9**: Representative western blots depicting analysis of co-immunoprecipitation studies performed to confirm the presence of multiple KRAS molecules in the native nanodiscs. Expi293 cells were transiently transfected with combinations of GFP-KRas and FLAG-KRas (+/- BI-2852). Native nanodiscs were purified using anti-FLAG M2 resin. Samples were blotted for Na/K ATPase (sample loading control), GFP, and FLAG using the following antibodies – anti- Na/K ATPase (Abcam ab167390), anti-GFP (CST 2956), and anti-FLAG (Sigma SAB4200071). Red asterisks denote the GFP-KRas bands eluted off of the anti-FLAG resin, which were subject to quantification by band densitometry using Fiji as summarized in *Fig. 4d*. All primary antibodies were used at 1:1000 dilution.

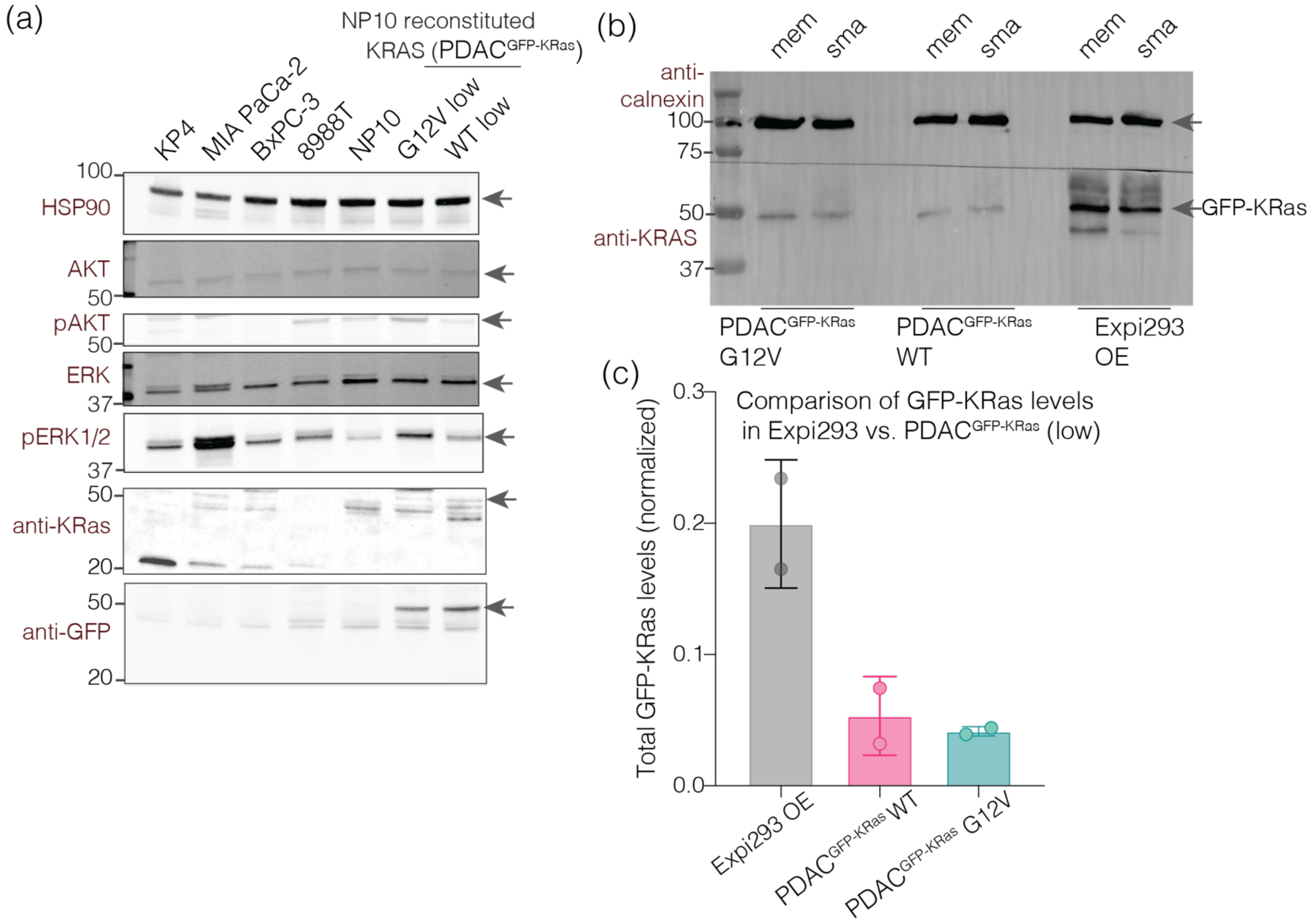

**Supplementary Figure 10**: KRas expression levels in various cell lines. (a) Western blot analysis comparing levels of various proteins in stable PDAC$^{GFP\text{-}KRas}$ cells expressing low levels of GFP-tagged KRas variants as compared to a panel of various PDAC cell lines expressing endogenous levels of KRas. Blots depict levels of a protein loading control (HSP90), endogenous and GFP-tagged KRAS (anti-KRAS and anti-GFP respectively), as well as the levels of important signaling molecules downstream of KRas (AKT and ERK), and the levels of their phosphorylated forms (pAKT, pERK1/2). (b) Representative western blot depicting the levels of GFP-KRas expressed in Expi293 cells and PDAC$^{GFP\text{-}KRas}$ cells stably expressing low levels of GFP-KRas or oncogenic KRas-G12V along with a loading control (calnexin). Lanes labelled 'mem' indicate isolated membrane fraction from respective cell lines were loaded. Lanes labelled 'sma' indicate that amphipathic copolymer-solubilized membranes (native nanodiscs) were loaded. (c) GFP-KRas expression levels in each cell line were quantified using band densitometry of the 'sma' labeled lanes and normalized with respect to total amounts of calnexin using Fiji. All primary antibodies were used at 1:1000 dilution. Data is shown as mean ± sd over three biological replicates.

### Supplementary Tables

**Table S1**: List of proteins used in this study with their number of transmembrane helices/subunit, oligomeric status, calculated molecular weights, and the position of the GFP-tag.

| Protein | # Transmembrane Domains/subunit | Oligomeric status | Molecular weight | Position of mEGFP tag | Expression cell lines |
|---|---|---|---|---|---|
| **SemiSWEET** | 3 | dimer | 10,816 Da | C-term | E. coli |
| **AmtB** | 11 | trimer | 44,515 Da | C-term | E. coli |
| **KcsA** | 2 | tetramer | 17,694 Da | C-term | E. coli |
| **LeuT** | 12 | Monomer:dimer | 57,408 Da | C-term | E. coli |
| **TrkA** | 1 | Monomer:dimer:higher order | 87,497 Da | C-term | SH-SY5Y, Expi293 |
| **KRas** | N/A (peripheral MP) | Monomer:dimer:higher order | 21,656 Da | N-term | Expi293, PDAC |

**Table S2**: Information for constructs obtained through Addgene.

| Plasmid name | Plasmid depositor | Addgene catalog # | Addgene link | Plasmid RRID | Plasmid reference |
|---|---|---|---|---|---|
| pEG BacMam N term His8 eGFP 3C | Dr. Eric Gouaux | 160684 | https://www.addgene.org/160684/ | Addgene_160684 | doi: 10.1038/nprot.2014.173 |
| pEG BacMam C term 3C eGFP His8 | Dr. Eric Gouaux | 160687 | https://www.addgene.org/160687/ | Addgene_160687 | doi: 10.1038/nprot.2014.173 |

**Table S3**: List of lysis, membrane resuspension, and FSEC buffers used in our experiments for different membrane proteins studied.

| | KRas | &TrkA | SemiSWEET, AmtB, LeuT WT and LeuT AA-mutant | KcsA |
|---|---|---|---|---|
| ***Lysis buffer** | 20 mM Tris-HCl pH 7.4, 300 mM NaCl or 20 mM HEPES pH 7.4, 300 mM NaCl | 25 mM Tris-HCl pH 8.0, 150 mM KCl | 50 mM Tris pH 7.4, 150 mM NaCl | 20 mM Tris-HCl pH 7.5, 100 mM KCl |
| ***,#Membrane resuspension buffer** | 50 mM Tris-HCl pH 8.1, 300 mM NaCl, 10% glycerol or 20 mM HEPES 7.4, 300 mM NaCl | 20 mM Tris-HCl, pH 8.0, 150 mM NaCl, 5 % glycerol | 50 mM Tris-HCl pH 8.1, 300 mM NaCl, 10% glycerol | 20 mM Tris-HCl pH 7.5, 100 mM KCl |
| **FSEC buffer** | 20 mM Tris pH 8.0, 150 mM NaCl or 20 mM | 20 mM Tris pH 8.0, 150 mM NaCl | 20 mM Tris pH 8.0, 150 mM NaCl | 20 mM Tris pH 7.5, 150 mM KCl |

| | |
|---|---|
| | HEPES pH 7.4, 150 mM NaCl |

*All lysis and membrane resuspension buffers were supplemented with PIC tablets (Pierce, Thermo).

#For membrane resuspension buffers, 1-2% of final concentration of copolymers (SMA or AASTY) were added.

&For TrkA, lysis buffer is also supplemented with phosphatase inhibitors.
